# Alzheimer’s disease alters astrocytic functions related to neuronal support and transcellular internalization of mitochondria

**DOI:** 10.1101/2021.09.15.460570

**Authors:** Riikka Lampinen, Irina Belaya, Liudmila Saveleva, Jeffrey R Liddell, Dzhessi Rait, Mikko T Huuskonen, Raisa Giniatullina, Annika Sorvari, Liisi Soppela, Nikita Mikhailov, Isabella Boccuni, Rashid Giniatullin, Marcela Cruz-Haces, Julia Konovalova, Marja Koskuvi, Tuomas Rauramaa, Andrii Domanskyi, Riikka H Hämäläinen, Gundars Goldsteins, Jari Koistinaho, Tarja Malm, Sweelin Chew, Kirsi Rilla, Anthony R White, Nicholas Marsh-Armstrong, Katja M Kanninen

## Abstract

Under physiological conditions *in vivo* astrocytes internalize and degrade neuronal mitochondria in a process called transmitophagy. Mitophagy is widely reported to be impaired in neurodegeneration but it is unknown whether and how transmitophagy is altered in Alzheimer’s disease (AD). Here we report that the internalization and degradation of neuronal mitochondria are significantly increased in astrocytes isolated from aged AD mouse brains. We also demonstrate for the first time a similar phenomenon between human neurons and AD astrocytes, and in murine hippocampi *in vivo*. The results suggest the involvement of S100a4 in impaired mitochondrial transfer between neurons and aged AD astrocytes. Significant increases in the mitophagy regulator Ambra1 were observed in the aged AD astrocytes. These findings demonstrate altered neuron-supporting functions of aged AD astrocytes and provide a starting point for studying the molecular mechanisms of transmitophagy in AD.

## 1. Introduction

Alzheimer’s Disease (AD) is a major cause of dementia, a progressive neurodegenerative disorder with increased risk upon aging. The pathology of the AD is typified by extracellular beta-amyloid plaques (Aβ), intracellular neurofibrillary tangles, neuronal loss, neuroinflammation and oxidative stress. However, despite extensive research, the actual causes of neurodegeneration remain unclear and there is no cure for the disease. (Lane et al., 2018)

The brain is an organ with an exceptionally high energy need. The mitochondria are central for energy metabolism, converting glucose to adenosine triphosphate (ATP) via oxidative phosphorylation. Glucose is considered to be the main energy source for neurons, and thus the brain is highly sensitive to changes in the mitochondrial function of cells. In addition, mitochondria are central for maintaining calcium homeostasis and various cell signaling pathways. Given that upon damage mitochondria release apoptotic factors such as cytochrome c, and that the mitochondria are a major source of reactive oxygen species (ROS), maintenance of healthy mitochondria is highly important for cellular well-being. Mitochondria quality is controlled by fusion and fission, intracellular localization, and permanent degradation of dysfunctional mitochondria via mitophagy. Impairments in mitophagy, selective form of autophagy, cause the intracellular accumulation of damaged mitochondria and are associated with adverse effects for the health of the brain cells. Cells with especially high risk are the long-lived neuronal cells. (Wang et al., 2020)

Mitochondrial dysfunction is associated with AD-induced neurodegeneration and aging. Synaptic mitochondria accumulate beta-amyloid (Aβ)_1-40_ and Aβ_1-42_, leading to impaired mitochondrial function and dynamics in neurons (Wang et al., 2016). A reduction in mitochondrial number and altered phenotypes are observed in the presynaptic regions of neurons of AD patients (Pickett et al., 2018). Given that neurons are post-mitotic cells, they also accumulate dysfunctional mitochondria upon aging. Furthermore, defective mitophagy is observed in AD-affected neurons (Ye et al., 2015) and AD patient brains have reported to contain reduced levels of autophagy-inducing protein beclin 1 (Pickford et al., 2008). Deficiency in beclin 1 have been shown to enhance Aβ deposition in mice modeling AD as well (Pickford et al., 2008).

The majority of studies deciphering aging- and AD-associated changes in mitochondria have in the past focused on neurons. Only relatively recently, astrocytes have started to gain larger interest due to their essential roles in maintaining brain health and implications in disease. Astrocyte reactivity is a common feature observed both in AD and upon aging (Clarke et al., 2018; Liddelow et al., 2017). Accumulation of Aβ_1-42_ causes adverse effects in astrocytes by impairing their mitochondrial function (Yao et al., 2018) and causing autophagy inhibition (Hong et al., 2018). However, to date, there are no reports assessing mitophagy in AD-affected astrocytes.

To meet the requirements for functional mitochondria, many cell types have been reported to be capable of transcellular mitochondrial movement. For example, transfer of mitochondria from mesenchymal stem cells to damaged or stressed cells of various cell types has been shown to be an important means of cellular regeneration and repair (Soundara Rajan et al., 2020). Tunneling nanotubes (TNTs) are suggested as one possible means of mitochondrial transfer between cells (Rustom et al., 2004; Soundara Rajan et al., 2020). Transmitophagy, the transfer of mitochondria to neighboring cells specifically for degradation, was termed for the first time by (Davis et al., 2014). The authors reported that degradation of retinal ganglion cell axon mitochondria occurs inside adjacent astrocytes under normal physiological conditions. Recently, mitophagy of dopaminergic neuron mitochondria was also suggested to be completed in neighboring astrocytes via spheroid-mediated transmitophagy. It was suggested that the aging of the astrocytes could lead to a failure in spheroid-mediated transmitophagy and in this way take part in the pathogenesis of Parkinson’s disease. (Morales et al., 2020)

Astrocytes are acknowledged to be critical for brain health, and to be affected by neurodegenerative diseases, but their functions related to mitochondria are less studied. The field is lacking detailed knowledge on the role of astrocytic mitophagy and transmitophagy in AD. Here we assessed how AD alters astrocytic functions related to neuronal support with a specific focus on mitochondria-related mechanisms, including transmitophagy. We were interested whether the AD-affected astrocytes are capable of internalizing and degrading neuronal mitochondria and whether the process is possibly altered due to AD-related changes in the basic functions of the astrocytes or their mitochondria.

## 2. Material and methods

### 2. 1. Animals

All experiments were approved by the National Animal Experiment Board of Finland and performed according to the animal protection guidelines of the Council of the European Union. The mouse lines used in this study were 5xFAD (on a C57BL/6J background) and C57BL/6J. The 5xFAD mice express human amyloid precursor protein (APP) and presenilin 1 (PS1) as transgenes with a total of five mutations causing familial AD (FAD) in these transgenes, three mutations in the *APP* (K670N/M671L, I716V, and V717I) and two in the *PS1* (M146L and L286V) (Oakley et al., 2006).

### 2.2. MitoEGFPmCherry and GFP lentiviral constructs

The plasmid for MitoEGFPmCherry was kindly provided by Professor Marsh-Armstrong, University of California, Davis (Davis et al., 2014). The plasmid was used to re-clone the transgenes to pCDH lentivirus transfer vector, in order to produce third generation lentiviral vectors. The production and purification of the lentiviral vectors were carried out at the laboratory of Docent Andrii Domanskyi, University of Helsinki. In the lentiviral vector the transgene expression was driven by the human synapsin (hSYN) gene promoter. The construct used in this study for astrocytes took advantage of a human phosphoglycerate kinase (hPGK) promoter to drive ubiquitous expression of green fluorescent protein (GFP) in the cells. This was produced by the Biocenter Kuopio Viral Gene Transfer Core.

### 2.3. Primary adult astrocyte cultures

The primary astrocytes were harvested from 2-3-, 5-6- and 10-12-month-old (mo) 5xFAD mouse brains as described in (Iram et al., 2016; Konttinen et al., 2019) with the following modifications. Briefly, the brains without cerebellum, olfactory bulb and brainstem were dissociated to single cell suspension with Adult Brain Dissociation kit (Miltenyi Biotech, Bergisch Gladbach, Germany) following the manufacturer’s instructions. The cells were cultured in DMEM/F-12 with GlutaMAX supplemented with 10% iFBS and 1% P/S (all Thermo Fisher Scientific, Waltham, MA, USA) on poly-D-lysine (Sigma-Aldrich, Saint Louis, MO, USA) coated 6-wells. The medium was changed daily for the 7 first days. The cells were split on day 7-8 *in vitro* (DIV) and on 10-11 DIV to expand the cultures. The cells were used for experiments at 14-26 DIV. 0.25% Trypsin–EDTA was used for detaching the cells. For studying internalization of neuronal mitochondria by astrocytes, the primary astrocytes were transduced with the lentivirus vector driving GFP expression in the cells with MOI 5 or 7 for 48 hours. The proportion of different cell types in the murine adult astrocyte cultures were assessed with qPCR and immunostaining for several markers of astrocytes, neurons, and microglia. The adult astrocyte cultures contain more than 80-85 % of astrocytes and a small proportion of other glial cells. Neurons were not detected in the cultures.

### 2.4. Primary cortical neuronal cultures

Primary cortical neuron cultures were prepared using the cortices of C57BL/6J mice on embryonic day 15 as described in (Loppi et al., 2021). The cells plated for experiments on poly-D-lysine coated plates (10 µg/ml in water, Sigma-Aldrich) in Neurobasal media supplemented with 2% B27, 1% penicillin-streptomycin (10,000 U/ml) (all Thermo Fisher Scientific) and 0.5 mM L-glutamine (Lonza, Walkersville, MD, USA). On 4 or 5 DIV, half of the media was replaced with fresh media to supply the cells with efficient amount of nutrients, unless the neurons were transduced or stained with mitochondria-targeted dye already on 2 DIV. The cells were used for experiments on 6-7 DIV.

### 2.5. Primary neuron-astrocyte co-cultures

Primary cortical neurons were plated on 24-well plates in Neurobasal media supplemented with 2% B27, 1% penicillin-streptomycin (10,000 U/ml) (all Thermo Fisher Scientific) and 0.5 mM L-glutamine (Lonza) with 200000 neurons/well for the MTT assay and 120000 neurons/well for live cell imaging with rhodamine 123 dye. Primary astrocytes were seeded on 24-well transwells with 0.4 µm pore size (Sarstedt AG & Co. Nümbrecht, Germany) with 5000 astrocytes/transwell for 1-2 days before transferring the transwells on top of the wells with cortical neurons. The two cell types were co-cultured for a total of 2 days before experiments were carried out. The cells were exposed to 250 µM glutamate (Sigma-Aldrich) for 24 hours before the MTT assay or live cell imaging with rhodamine 123 dye.

For assessing the internalization and degradation of neuronal mitochondria by astrocytes, the primary neurons were plated in µ-Slide 8 wells at 100 000 neurons/well (ibidi GmbH, Gräfelfing, Germany). At 2 DIV, the neuronal cells were either transduced with the LV-mito-EGFP-mCherry-msSODUTR with MOI 1 for 24 hours or stained for 30 minutes at +37 °C with 100 nM MitoTracker Red CMXROS dye (Thermo Fisher Scientific). Three days after labeling the neuronal mitochondria either with the lentivector or MitoTracker, the primary astrocytes were seeded in co-cultures with the neurons. The cells were grown in co-cultures for 48 hours before fixing the cells with 4% formaldehyde.

### 2.6. iPSC-derived neuron- and astrocyte-like cultures

The induced pluripotent stem cells (iPSCs) used in this study were derived from control subjects described previously (Tiihonen et al., 2019). The iPSC lines were generated and differentiated to astroglial cells and neuronal cells using previously described protocols (Oksanen et al., 2017; Tiihonen et al., 2019). On day 4, after plating the neuronal cells as 75 000 cells/cm^2^ onto poly-ornithinine/Matrigel-coated glass coverslips, the neuronal cells were transduced with the LV-mito-EGFP-mCherry-msSODUTR at MOI 1 for 24 hours. The astrocyte progenitor cells derived from spheres were further maturated for one week in astrodifferentiation medium (DMEM/F12 supplemented with 1% N_2_ supplement, 1% Glutamax, 1% non-essential amino acids, 0.5% penicillin/streptomycin (50 IU/50 μg/mL), 0.5 IU/mL heparin (LEO Pharma, Ballerup, Denmark), 10 ng/mL bFGF and 10 ng/mL EGF (both growth factors from PeproTech EC Ltd., London, UK) prior to plating in co-culture with neurons. Following maturation, the astrocytes were detached with Accutase (STEMCELL Technologies, Vancouver, Canada) and replated in neural sphere medium at 10 000 astrocytes/cm^2^ on top of the lentivector-transduced neurons. Cells were grown as mixed cultures for 7 days prior to fixing the cultures with 4% paraformaldehyde.

### 2.7. Intracerebral injections of MitoEGFPmCherry lentivirus

*In vivo*, the localization of neuronal mitochondria inside 5xFAD mouse astrocytes was assessed following intracerebral injection of the LV-mito-EGFP-mCherry-msSODUTR to the hippocampi of 6 mo 5xFAD mice. The vector was injected in the volume of 2.5 µl into the dentate gyrus region of the hippocampus by using the following coordinates: ±3.2 mm medial/lateral, −2.7 mm anterior/posterior, −2.7 mm dorsal/ventral from the bregma as described previously (Kanninen et al., 2009). One week after the intracerebral injection with the viral vector, the mice were deeply anesthetized, perfused transcardially with heparinized saline and the brains were immersion-fixed in 4 % PFA for 22h as described previously (Kanninen et al., 2009). Following cryoprotection in sucrose solution, the brain tissues were frozen in liquid nitrogen and cut as 20 µm sections with cryostat (Leica Microsystems GmH, Wetzlar, Germany).

### 2.8. Immunocytochemistry

The co-cultures of lentivector-transduced murine primary neurons and astrocytes were fixed with 4% formaldehyde in DPBS for 20 minutes, permeabilized with 0.2% Triton X-100 in DPBS for 30 minutes and the nonspecific binding of antibodies was blocked with incubation with 5% normal goat serum in DPBS for 30 minutes at room temperature. For immunostaining of the astrocytes, the co-cultures were incubated first with primary antibody for glial fibrillary acidic protein (GFAP, 1:400, Z033429-2, Dako, Glostrup, Denmark) prepared in 5% NGS in DPBS overnight at +4 °C following an incubation with Alexa Fluor405 (A31556, Thermo Fisher Scientific, 1:500) or Alexa Fluor680 (A31556, Thermo Fisher Scientific, 1:2000) secondary antibodies prepared in 5% NGS in DPBS for 2 h at room temperature. For iPSC-derived neuron- and astrocyte-like cultures, the protocol for immunocytochemistry was similar with only minor modifications. The co-cultures were fixed with 4% PFA, permeabilized with 0.25% Triton X-100 in DPBS for 1h at room temperature and the blocking with 5% normal goat serum was extended for 1h. The primary antibody against GFAP (Dako, Z033429-2) was used at 1:500 dilution and Alexa Fluor405 secondary antibody at 1:500 dilution. The coverslips were mounted on glass slides with Vectashield mounting medium (Vector Laboratories INC, Burlingame, CA, USA) for fluorescence with 4′,6-diamidino-2-phenylindole (DAPI).

For visualizing tunneling nanotubes, the co-cultures of MitoTracker CMXROS labeled primary E15 neurons and astrocytes derived from adult WT or 5xFAD mouse brain were first fixed with 4% formaldehyde and then labelled with Alexa Fluor 488 Phalloidin dye (Thermo Fisher Scientific) according to the manufacturer’s instructions. Co-cultures were imaged with a Zeiss Axio Observer inverted microscope with LSM800 confocal module with 63x objective and ZEN software v.2.3 (Carl Zeiss AG, Oberkochen, Germany). For the experiments studying the induction of TNT-like structures with H_2_O_2_, the neurons were treated with 1 µM H_2_O_2_ for 2 h prior labeling the neurons with Mitotracker CMXROS dye.

### 2.9. Immunohistochemistry

Mouse brain cryosections for studying transmitophagy *in vivo* were blocked with 10% normal goat serum for 30 minutes at room temperature and incubated with an anti-GFAP primary antibody (Dako, Z033429-2, 1:500 dilution in 5% normal goat serum) overnight at room temperature. Next, the sections were washed with PBS containing 0.2% Tween20 (Sigma-Aldrich) and incubated with Alexa Fluor 405 (A31556, Thermo Fisher Scientific, 1:500 dilution in 5% normal goat serum) for 2 h at room temperature. After washes with PBS containing 0.2% Tween20 the sections were embedded with Fluoromount-G mounting medium (SouthernBiotech, Birmingham, AL, USA).

To study neuronal localization of S100a4 in the hippocampal area of 12 mo WT and 5xFAD mice, the mice were deeply anesthetized and transcardially perfused as described above. The brains were post-fixed in 4% PFA for 21h at +4 °C, cryoprotected with immersion to 30% sucrose for 48h at +4 °C and finally frozen at -70 °C prior sectioning. The 20 µm sagittal sections of one hemisphere were cut with 400 µm interval. The sections were rehydrated overnight in 0.1 M PB and washed with 1xPBS before antigen retrieval with boiling the sections in 10mM citrate. Endogenous peroxidases were blocked by treating sections with 0.3% H_2_O_2_ in methanol for 30 minutes. Next, the sections were blocked with 0.5% Mouse on Mouse Blocking Reagent (Vector Laboratories, MKB-2213) for 1h at RT with following blocking with TSA blocking reagent (Perkin Elmer, Waltham, MA, USA, FP1020) as 0.5% solution in PBS pH 7.4 for another 1h at RT. Sections were incubated o/n at RT with primary antibodies diluted in 0.5% solution of the TSA blocking reagent. Primary antibodies used for assessing the neuronal localization of S100a4 were anti-NeuN (MAB377, Sigma-Aldrich, dilution 1:200) and anti-S100a4 (ab41532, Abcam, Cambridge, UK, dilution 1:100). Incubation with the secondary antibodies was performed for 2h at RT. Secondary antibodies biotin conjugated goat anti-rabbit (BA-1000, Vector Laboratories, dilution 1:200) and goat anti-mouse IgG (H+L) Alexa Fluor 488 (A11001, Thermo Fisher Scientific, dilution 1:250) were diluted in 0.5% solution of the TSA blocking reagent. Sections were further processed according to the instructions in the TSA Plus Cyanine 3 kit (Perkin Elmer, NEL744001KT) in order to visualize the biotin conjugated secondary antibody bound to anti-S100a4 primary antibody. Lastly, the sections were embedded with Vectashield mounting medium (Vector Laboratories) for fluorescence with 4′,6-diamidino-2-phenylindole (DAPI). For visualizing S100a4 together with astrocytic marker GFAP and amyloid plaques, same protocol was used with anti-S100a4 (ab41532, Abcam, dilution 1:100), anti-GFAP (ab4674, Abcam, dilution 1:2500) and Anti-Amyloid β, clone W0-2 (MABN10, Sigma-Aldrich, dilution 1:1000) primary antibodies. Secondary antibodies used in this staining were biotin conjugated goat anti-rabbit (BA-1000, Vector Laboratories, dilution 1:200), goat anti-chicken IgY (H+L) Alexa Fluor 488 (A11039, Thermo Fisher Scientific, dilution 1:250) and goat anti-mouse IgG (H+L) Alexa Fluor 680 (A21057, Thermo Fisher Scientific, dilution 1:250).

To study neuronal localization of S100a4 in the human brain, paraffin blocks from posterior hippocampus were cut to 5 µm sections and anti-S100a4 immunostaining was combined with Nissl-staining. There were two samples from patients with Alzheimer’s disease and two control cases in the staining. The study was approved by the Ethics Committee of the Hospital District of Northern Savonia (276/2016). First, the paraffin sections were deparaffinized using xylene and rehydrated with decreasing concentrations of ethanol. Next, the sections were washed with 1xPBS before antigen retrieval with boiling the sections in 10 mM citrate. Endogenous peroxidases were blocked by treating sections with 0.3% H_2_O_2_ in methanol for 30 minutes and followed with blocking with TSA blocking reagent (Perkin Elmer) as 0.5% solution in PBS pH 7.4 for 1 h at RT. Sections were incubated o/n at RT with primary antibodies diluted in 0.5% solution of the TSA blocking reagent. The primary antibody used was anti-S100a4 (ab41532, Abcam, dilution 1:100). Incubation with the secondary antibody (Biotin conjugated goat anti-rabbit (BA-1000, Vector Laboratories, dilution 1:200)) diluted in 0.5% solution of the TSA blocking reagent was performed for 2 h at RT. Sections were further processed according to the instructions in the TSA Plus Cyanine 3 kit (Perkin Elmer, NEL744001KT) to visualize the biotin conjugated secondary antibody bound to anti-S100a4 primary antibody. The sections were imaged for representative images without letting the sections to dry with Zeiss Axio Imager 2 fluorescent microscope with 10x objective before continuing the Nissl-staining. The Nissl-staining was performed by first rinsing the sections with water, followed with incubation with solution of thionin acetate, after which the sections were washed twice with water and once with both 50 % and 70 % ethanol. Finally, the sections were mounted with xylene before imaging for Nissl for the same locations as the images for anti-S100a4 staining were taken previously.

Stained sections were imaged with Zeiss Axio Imager 2 fluorescent microscope with 10x objective for all the immunohistochemical staining’s of S100a4 and ZEN software (Carl Zeiss AG). Images were analyzed with ImageJ for quantification of percentage of immunoreactive area for S100a4 in selected regions.

### 2.10. Assessing the internalization and transmitophagy of neuronal mitochondria by astrocytes *in vitro* and *in vivo*

Transmitophagy and internalization of the lentivirus labelled neuronal mitochondria *in vitro* and *in vivo* were visualized by imaging with a Zeiss Axio Observer inverted microscope with LSM800 confocal module with 63x objective and ZEN software v. 2.3 (Carl Zeiss AG, Oberkochen, Germany). For quantifying the internalized and degraded neuronal mitochondria in primary murine or iPSC-derived astrocytes, the number of intact and degraded neuronal mitochondria were manually counted per one astrocyte from confocal z-stack images aided with the profile tool in the ZEN software. The results were counted as an average of three biologically individual experiments for each age-group for murine astrocytes and as per one donor for iPSC-astrocytes.

### 2.11. Measurement of neuronal metabolic activity

Metabolic activity of the neurons co-cultured with astrocytes isolated from 5-6 mo and 11-12 mo WT or 5xFAD mice was assessed with the MTT assay. Briefly, the cell culture medium was replaced with fresh cell culture medium supplemented with 1.2 mM MTT ((3-(4, 5-dimethylthiazolyl-2)-2, 5-diphenyltetrazolium bromide), Sigma-Aldrich). For lysed cell control the cells we treated with 30% (v/v) Triton X-100 (Sigma-Aldrich) for 5 minutes prior replacing the medium. The cells were incubated in the media containing MTT for 1-4 hours at +37 °C before solubilizing the cells with dimethyl sulfoxide. Absorbance of 100 µl aliquots of solubilized cells was measured at 595 nm with Wallac Victor 1420 microplate reader (Perkin Elmer).

### 2.12. Measuring mitochondrial membrane potential with rhodamine123 imaging

Cell cultures (astrocytes or neurons) were incubated for 30 min at 37 °C in 5 µM rhodamine123 solution (ThermoFisher Scientific, 5 mM stock solution in 99% EtOH). Then cells were transferred to the imaging system where they were constantly perfused with Basic Salt Solution (BSS, contained in mM 152 NaCl, 10 HEPES, 10 glucose, 2.5 mM KCl, 2 CaCl_2_, 1 MgCl_2_. pH was adjusted to 7.4). First, the baseline was recorded for 1 min before applying 4 µM FCCP (Abcam, 20 mM stock solution in DMSO) for 2 min. The response was calculated as ΔF/F_0_ (normalized to baseline). Both BSS and FCCP solution contained 0.02% (v/v) DMSO. Our TILL Photonics imaging system (TILL Photonics GmbH, Germany) was equipped with fast perfusion system (Rapid Solution Changer RSC-200, BioLogic Science Instruments, Seyssinet-Pariset, France), allowing fast exchange between applying solutions (∼ 30 ms). Cells were imaged with Olympus IX-70 (Olympus Corporation, Tokyo, Japan) with CCD camera (SensiCam, PCO imaging, Kehlheim, Germany) with 10x objective for astrocytes or 20x objective for neurons. The excitation wavelength was 495 nm. Imaging was conducted at 1 FPS. All the experiments were conducted using Live Acquisition and processed with Offline Analysis software (TILL Photonics GmbH, Munich, Germany).

### 2.13. Western blotting

Cells were lysed directly in 1x Laemmli buffer (62.5 mM Tris-HCl (pH 6.8), 2.3% SDS, 5% β-mercaptoethanol, 10% glycerol, 0.02% bromophenol blue). The cell lysates were boiled at 95 °C and run on 10% or 15% SDS-PAGE Tris-glycine gels. The proteins were transferred to PVDF membranes with Trans-Blot Turbo Transfer System (Bio-Rad, Hercules, CA, USA) following blocking with 5% non-fat dry milk solution prepared in 0.2 % Tween-20/0.01 M PBS for 30 min. The immunodetection of selected proteins was performed overnight at 4 °C with the following antibodies: p62 (Cell Signaling Technology Inc, Danvers, MA, USA, 5114, 1:1000), LC3b (Abcam ab51520, 1:3000 dilution), Tom20 (Proteintech Group Inc, Rosemont, IL, USA, 11802-1-AP, 1:2000 dilution), Ambra1 (Proteintech 13762-1-AP, 1:200 dilution) and β-actin (Sigma-Aldrich A5441, 1:5000) for loading control. For detection of the proteins of interest, the membranes were incubated for 2 h at room temperature in HRP conjugated IgG anti-rabbit secondary antibody (Bio-Rad 170-65-15, 1:3000) or Cy5 conjugated IgG anti-mouse secondary antibody (Jackson ImmunoResearch Laboratories Europe Ltd., Cambridgeshire, UK 715-175-151, 1:1000 dilution). The membranes incubated with HRP conjugated secondary antibody were further developed using enhanced chemiluminescence (SuperSignal™ West Pico PLUS Chemiluminescent Substrate, Thermo Fisher Scientific). All membranes were imaged on a Bio-Rad ChemiDoc XRS+ System.

### 2.14. Statistical analyses and graphical illustrations

The data was analyzed using t-test or ANOVA as appropriate using GraphPad Prism 8.1.0 (GraphPad Software Inc, San Diego, CA, USA). Before performing the statistical test, the data was analyzed for normality and possible outliers were identified with the ROUT method (Q=1%) in GraphPad Prism. Statistical significance was assumed if P < 0.05 and confidence intervals were reported with SEM. The graphical illustrations were created with BioRender.com.

Material and methods for supplementary figures and tables are described in separate supplementary file.

## 3. Results

### 3.1. Transcellular internalization and degradation of neuronal mitochondria is altered in aged AD astrocytes

Movement of mitochondria between neurons and astrocytes has been shown to occur in normal physiological conditions in the mouse optic nerve head *in vivo* (Davis et al., 2014), after stroke from astrocytes to neurons both *in vitro* and *in vivo* (Hayakawa et al., 2016) and to rescue cisplatin-treated neurons (English et al., 2020). Here we utilized two fluorescent mitochondrial labels to study whether astrocytes internalize mitochondria derived from neurons, and whether the process is altered in AD. Neuronal mitochondria were labelled with a MitoTracker CMXROS dye, or a mitochondria-targeted tandem fluorophore reporter before co-culturing neurons with astrocytes. Confocal imaging revealed the presence of neuron-derived mitochondria inside both WT and AD astrocytes (Fig 1A and 1B). Astrocytes harvested from 5xFAD mouse brains internalized significantly more neuronal mitochondria than their WT controls in all age groups. The aged (11 mo) 5xFAD astrocytes internalized the most neuronal mitochondria (mean difference 27.13 ± 6.805, p=0.0002) compared to the wild-type (WT) astrocytes (Fig 1C). The increased internalization of the neuronal mitochondria by AD astrocytes was also confirmed with human-derived cells. The iPSC-astrocytes derived from a symptomatic AD patient with the PSEN1 ΔE9 mutation were observed to internalize significantly more neuronal mitochondria derived from iPSC-neurons compared to its isogenic, PSEN1 mutation corrected, control cells (mean difference 4,960 ± 1,554, p=0.0023). Interestingly, there was no significant difference in internalization of the neuronal mitochondria between the iPSC-astrocytes derived from pre-symptomatic AD patient and its isogenic control cells (Fig 1E). Thin nanotube-like structures consisting of filamentous actin were observed to be bridging between the primary neurons and astrocytes in the cultures (Fig. 1F), suggesting one possible route allowing the transcellular movement of mitochondria from neurons to astrocytes. The H_2_O_2_ treatment on neurons seemed to increase the thin TNT-like protrusions around astrocytes, and even more in 5xFAD astrocytes compared to WT astrocytes (Fig 1G).

**Figure 1.**
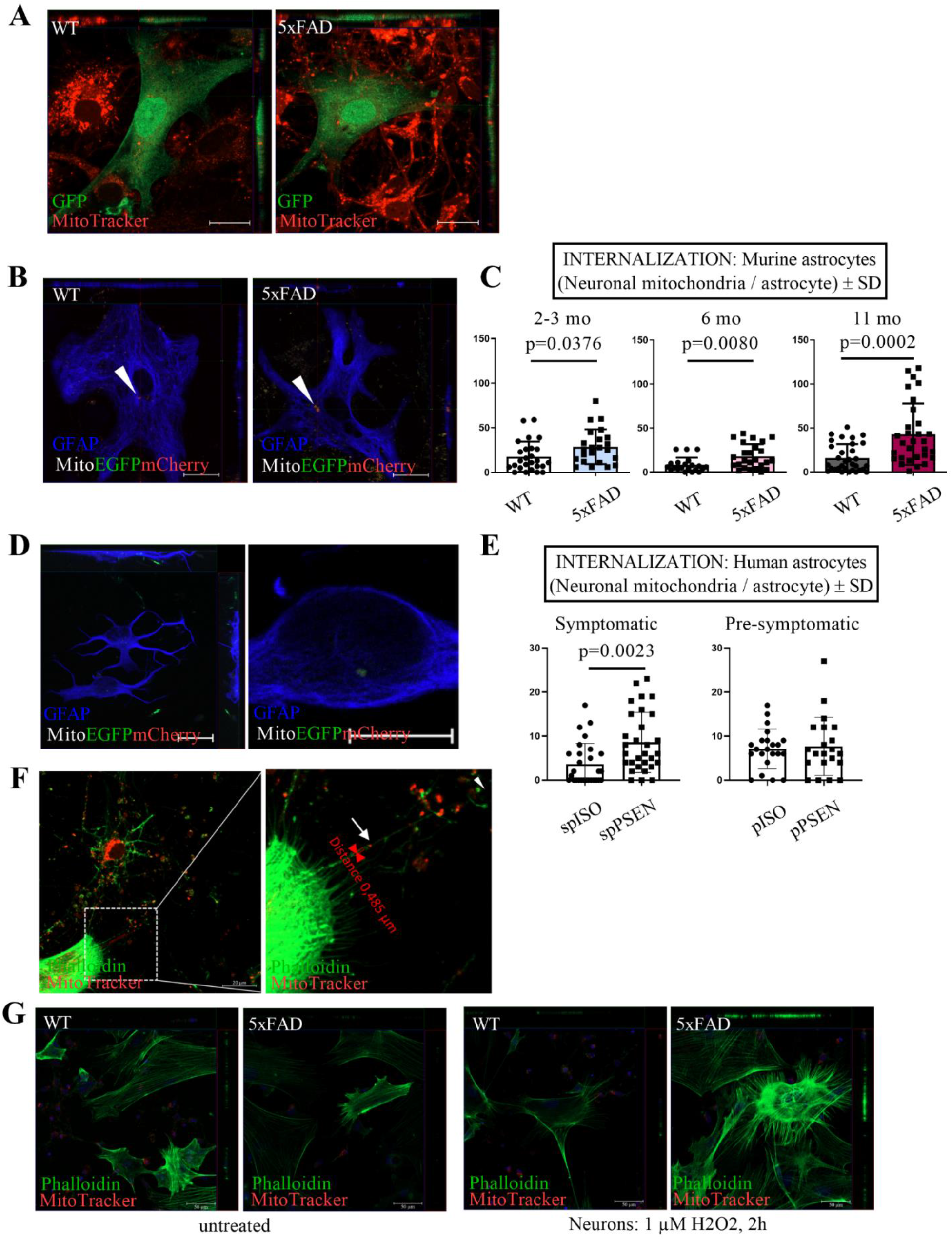
Astrocytes’ ability to internalize neuronal mitochondria is altered in AD. (A) Example confocal images from one z-plane showing neuronal mitochondria internalized by astrocytes *in vitro* in orthogonal view. E15 cortical neurons were labelled with MitoTracker CMXROS dye prior to co-culturing with adult astrocytes derived from 5 mo WT or 5xFAD mouse brains and expressing a lentiviral-GFP construct. Sacle bar 20 µm. (B) Example confocal images from one z-plane showing internalized neuronal mitochondria inside adult astrocytes *in vitro* in orthogonal view. Adult astrocytes were co-cultured with E15 neurons expressing a lentiviral-mitoEGFPmCherry construct. Scale bar 20 µm. (C) Quantified amounts of internalized neuronal mitochondria in astrocytes harvested from 2-3 mo, 6 mo and 11 mo WT and 5xFAD mouse brain. Neuronal mitochondria were labeled with a lentiviral-mitoEGFPmCherry construct. Data is shown as all (mCherry only and mCherry + EGFP) fluorescence signal peaks/cell ± SD. N=3 biologically independent replicates for each age-point. Each dot represents one single astrocyte imaged. Three biologically independent experiments were carried out for each age-point. Unpaired two-tailed t test. (D) Example images showing degraded neuronal mitochondria inside astrocyte in iPSC-derived neuron- and astrocyte cultures. An internalized mitochondria is shown in maximum intensity projection of a z-stack (left hand side, scale bar 20 µm) and as a digital zoom-in from one z-stack plane for an iPSC-astrocyte with internalized neuronal mitochondria (right hand side, scale bar 10 µm).(E) Quantified amounts of neuronal mitochondria in iPSC-astrocytes derived from symptomatic and pre-symptomatic AD patients carrying PSEN1 mutation and their isogenic (mutation corrected) lines. N=22-30 analyzed astrocytes/ iPSC-line. spPSEN, symptomatic donor with clinical diagnosis for AD and PSEN1 ΔE9 mutation. pPSEN, pre-symptomatic donor with PSEN1 ΔE9 mutation. SpISO, isogenic (PSEN1 mutation corrected) line for the symptomatic donor, pISO, isogenic (PSEN1 mutation corrected) line for the pre-symptomatic donor. Unpaired two-tailed t-test. (F) An example confocal image showing neuronal mitochondria traveling along a tunneling nanotube visualized with phalloidin staining. Neuronal mitochondria were labelled with MitoTracker CMXROS dye prior co-culturing with adult astrocytes harvested from 3 mo 5xFAD mouse brain. On the right-hand side an enlarged image of the boxed area. The white arrow heads point to examples of degraded neuronal mitochondria in astrocytes and white arrow point to neuronal mitochondria traveling along the TNT-like structure. Scale bar 20 µm. Example images acquired with objective with 63x objective. (G) Example images of murine neuron-astrocyte co-cultures where formation of TNT-like structures were induced in astrocytes by treating the neurons with 1 µM H2O2 for 2h prior labeling the neuronal mitochondria with Mitotracker CMXROS dye and co-culturing them with astrocytes. TNT-like structures were visualized with phalloidin dye. Scale bar 50 µm. All graphs represent the mean ± SD.

To validate whether astrocytes can degrade neuron-derived mitochondria *in vitro*, we transduced primary neurons with a tandem fluorophore reporter of acidified mitochondria (Fig 2A) before co-culturing the neurons with astrocytes. Confocal imaging revealed the presence of neuron-derived, degraded mitochondria inside the cultured astrocytes (Fig 1B). Transmitophagy was also confirmed in human induced pluripotent stem cell (iPSC) derived astrocytes co-cultured with iPSC-derived neurons (Fig 1D) and in the mouse hippocampi *in vivo* (Fig 2D).

**Figure 2.**
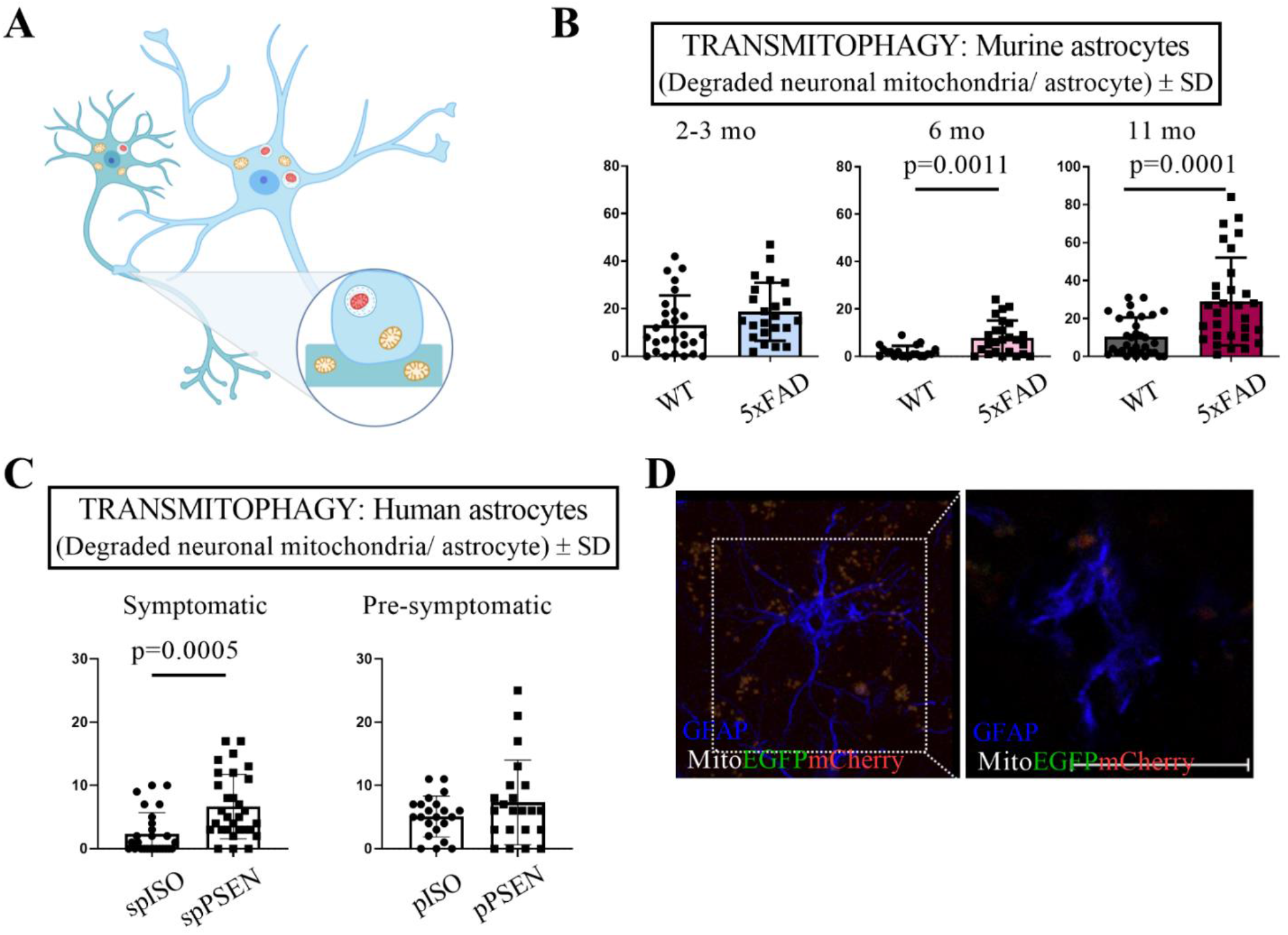
Astrocytes’ ability to degrade neuronal mitochondria is altered in aging and in AD. (A) Graphical illustration of the lentiviral-mitoEGFPmCherry construct. The synapsin promoter driven mitochondria-targeted reporter construct contains EGFP (green) and mCherry (red), which colocalize in mitochondria (yellow signal). Upon meeting the acidic environment of the lysosome, the EGFP signal is lost, resulting in red mCherry fluorescence, indicative of mitochondrial degradation. (B) Quantified amounts of degraded neuronal mitochondria in astrocytes harvested from 2-3 mo, 6 mo and 10-11 mo WT and 5xFAD mouse brains. Neuronal mitochondria were labeled with lentiviral-mitoEGFPmCherry construct. Data is shown as only mCherry fluorescence signal peaks/cell ± SD. N=3 biologically independent replicates for each age-point. Each dot represents one single astrocyte imaged. Three biologically independent experiments were carried out for each age-point. Unpaired two-tailed t test. (C) Quantified amounts of degraded neuronal mitochondria in iPSC-astrocytes derived from symptomatic and pre-symptomatic AD patients carrying PSEN1 mutation and their isogenic (mutation corrected) lines. N=22-30 analyzed astrocytes/ iPSC-line. spPSEN, symptomatic donor with clinical diagnosis for AD and PSEN1 ΔE9 mutation. pPSEN, pre-symptomatic donor with PSEN1 ΔE9 mutation. SpISO, isogenic (PSEN1 mutation corrected) line for the symptomatic donor, pISO, isogenic (PSEN1 mutation corrected) line for the pre-symptomatic donor. Unpaired two-tailed t-test. (D) Maximum intensity projection of a z-stack and orthogonal view from one z-stack plane (right hand side) visualizing degraded neuronal mitochondria inside an astrocyte in the hippocampus of a 6 mo 5xFAD mouse and a digital zoom-in from one z-stack plane with internalized neuronal mitochondria (right hand side, scale bar 20 µm).

Impairments in mitochondrial quality control have previously been linked to AD (Lampinen et al., 2018; Reddy and Oliver, 2019) and AD-affected neurons have previously been reported to accumulate dysfunctional mitochondria (Fang et al., 2019). We next questioned whether transmitophagy alterations occur in AD. Co-cultures of WT neurons with astrocytes harvested from mouse brains at various ages demonstrated that the degree of transmitophagy was not changed in young (2-3 mo) astrocytes. However, AD astrocytes derived from both 6 mo and 11 mo mice displayed an increase (in the total amount of degradation of neuronal mitochondria (Fig 2B). Furthermore, similarly to the murine AD astrocytes, the iPSC-astrocytes derived from a symptomatic AD patient with the PSEN1 ΔE9 mutation were observed to degrade significantly more neuronal mitochondria derived from iPSC-neurons compared to its isogenic, PSEN1 mutation corrected, control cells (mean difference 4,296 ± 1,156, p=0.0005). However, no significant difference in degradation of the neuronal mitochondria between the iPSC-astrocytes derived from pre-symptomatic AD patient and its isogenic control cells was observed (Fig 2C). The degradation of neuronal mitochondria was the highest in murine astrocytes derived from 11 mo 5xFAD mice (mean difference 18,56 ± 4,478, p=0.0001) indicating an age-dependent alteration to transmitophagy in AD astrocytes.

### 3.2. Aged 5xFAD mice display a shift in immunoreactivity for S100a4 from neurons to astrocytes

The concentration gradient of the S100a4 protein between neurons and astrocytes has been shown to determine the direction of TNT formation, which serve as potential transcellular highways for mitochondrial transfer between these cell types (Sun et al., 2012). To determine its involvement in the mitochondrial transfer from neurons to astrocytes, we assessed levels of S100a4 by immunohistochemistry in 5xFAD mouse brain sections. The immunoreactivity of S100a4 was not significantly reduced in the areas highly enriched with hippocampal neurons of 12 mo 5xFAD mice when compared to age-matched WT mice, although there was seen a trend for reduced immunoreactivity in areas enriched with neurons the images of the anti-S100a4 immunostained brain sections (data not shown). On the other hand, in the fiber tract area, populated primarily by astrocytes, a significant increase (mean difference above CA1 4.300 ± 0.4966, p<0.0001 and mean difference above CA3 2.967 ± 0.3850, p<0.0001) in immunoreactivity for S100a4 was observed in 12 mo 5xFAD mice (Fig 3A). In addition, in 5xFAD sections the S100a4 staining was observed close to areas stained with anti-amyloid β antibody (Fig 3A).

**Figure 3.**
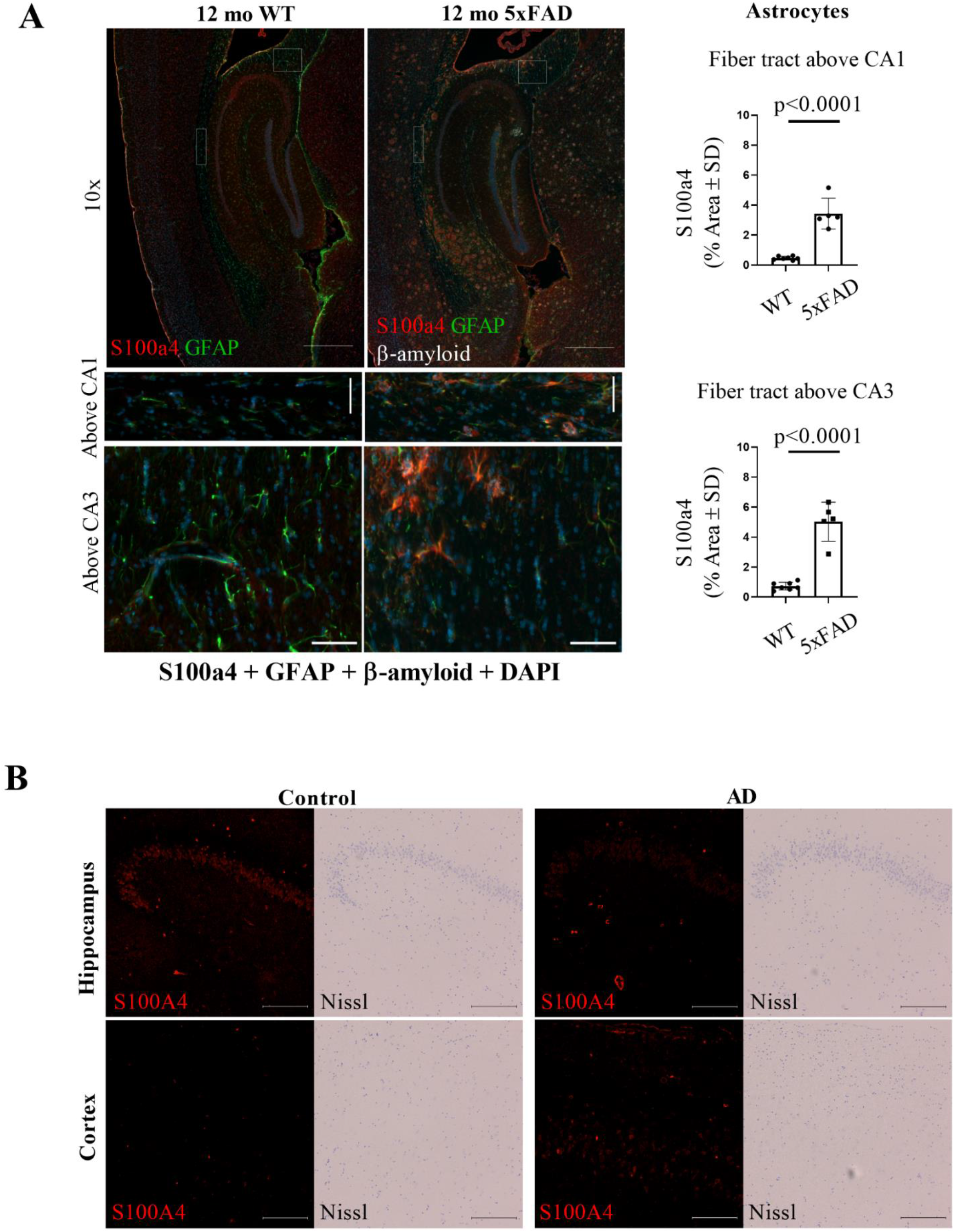
S100a4 is increased in aged 5xFAD astrocytes *in vivo*. (A) The immunoreactivity of S100a4 was by histochemical staining from cryosections of 12 mo 5xFAD and WT brain sections in cells resembling astrocytes by morphology in the fiber tract region above CA1 and CA3 neuronal layers. Quantitative data is presented as immunoreactive area for S100a4, % ± SD. N=7 WT mice and N=4 5xFAD mice. Example images with 10x objective (scale bar 500 µm) and enlargement of boxed areas (scale bar 50 µm) showing the difference between genotypes. Anti-S100a4 staining is shown as red, anti-GFAP as green, DAPI as blue and anti-amyloid β with white pseudo color. Unpaired two-tailed t test. (B) The immunoreactivity for S100a4 in neuron-enriched areas was assessed by immunohistochemical staining in paraffin sections of human brain posterior hippocampus. The location of neuronal cells was visualized with Nissl staining. Example images were taken with 10x objective. Scale bar 200 µm.

We performed further an ELISA assay for S100a4 from cell lysates of adult astrocytes derived from 5-6 mo and 11-12 mo WT and 5xFAD mice (Supplementary figure 1). The results indicate an increase of intracellular S100a4 in astrocytes harvested from 12 mo 5xFAD mice compared to the other study groups, similarly as immunostaining for S100a4 was observed to be increased in 12mo 5xFAD mouse brain in astrocyte-rich areas.

To assess the S100a4 immunoreactivity also in the human brain, we immunostained *post mortem* human brain sections with both anti-S100a4 antibody and Nissl staining. Based on these example images it seems that in the human brain the AD neurons have less S100A4 in the hippocampal layer, where the neurons are visualized with Nissl-staining. However, in the cortex the AD brain seemed to have more positive immunostaining for S100A4 in other cell types besides neurons (Fig 3B). These data are in line with our findings in the 5xFAD mice for the glial-like cells.

### 3.3. Mitochondrial functions of astrocytes, astrocyte reactivity and phagocytosis are unaltered upon aging and AD

The observation of increased internalization and degradation of neuronal mitochondria by aged 5xFAD astrocytes led us to investigate the functionality and health of the astrocytes’ own mitochondria. We first assessed cytochrome c oxidase (mitochondrial electron transport chain complex IV) activity in astrocytes in WT and 5xFAD mouse brains by histochemistry. The cytochrome c oxidase activity appeared to be altered specifically around beta-amyloid plaques in aged 5xFAD brains (Suppl. Fig. 2A), indicating that the mitochondrial function of a subset of astrocytes may be altered *in vivo*. However, we did not observe a difference in the basal respiration rate between age-matched 5xFAD and WT astrocytes (Suppl. Fig 2B). Upon comparing the 5xFAD astrocytes harvested from 5-6 mo mice to those extracted from 10-12 mo 5xFAD mice, the 5xFAD astrocytes from 5-6 mo mice was observed to have significantly higher basal respiration rate. Furthermore, levels of intracellular ATP were not altered when comparing WT and 5xFAD astrocytes (Suppl. Fig 2C) and live cell imaging of the mitochondrial membrane potential of astrocytes did not reveal alteration in 5xFAD astrocytes (Suppl. Fig. 2D). The absence of mitochondrial alterations in 5xFAD astrocytes was further supported by measurement of the mitochondrial content in the cells by Western blotting for the mitochondrial outer membrane protein Tom20, which remained unchanged during aging and in 5xFAD cells (Suppl. Fig 2E). To rule out the possibility of increased levels of oxidative stress in the aged 5xFAD astrocytes, the gene expression levels of the antioxidant-response genes heme oxygenase-1 (*Hmox1*) and NAD(P)H: quinone oxidoreductase (*Nqo1*) were assessed by qPCR. However, no significant difference between WT and 5xFAD astrocytes was observed (data not shown).

In rodent brains, the astrocytes have been reported to manifest as the reactive sub-phenotype, termed A1, in AD and normal aging (Clarke et al., 2018; Liddelow et al., 2017). To determine how aging and AD affect key astrocytic functions besides mitochondrial internalization and degradation, we assessed their reactive phenotype and phagocytic ability. The expression level of the A1 marker gene *Serping1* was observed to be increased and while the other A1 marker gene *Srgn* expression was reduced in astrocytes isolated from the adult 5xFAD mouse brains, indicating that the FAD mutations alter the phenotype of astrocytes (Suppl. Fig. 3A). The expression of genes encoding phagocytosis-related proteins was unchanged in AD and during aging of astrocytes except for up-regulation of *Megf10* (Log2 FC 1.0689, p<0.045) in 5xFAD astrocytes at 2-3 mo age compared to the astrocytes harvested from age-matched WT mice (Suppl. Fig. 3B). Furthermore, no difference was observed in phagocytosis of pHrodo Zymosan A bioparticles between the WT and 5xFAD astrocytes at all ages, indicating that the phagocytic capacity of astrocytes remains unaltered (Suppl. Fig 3C). The 5xFAD astrocytes were not observed to increase secretion of inflammatory cytokines compared to WT (Supplementary Tables 2 and 3).

### 3.4. Astrocyte-mediated neuronal support is altered in AD

We next assessed whether the ability of astrocytes to support neuronal functions is altered upon aging and/or AD, possibly thereby elucidating why the internalization and degradation of neuronal mitochondria is increased in the aged 5xFAD astrocytes. Based on live cell imaging with rhodamine 123 dye, co-culturing primary neurons with 5xFAD astrocytes from aged mice altered the mitochondrial membrane potential (MMP) of neurons (Fig 4A). When the neurons were co-cultured with 5xFAD astrocytes and treated with glutamate (mimicking the glutamate excitotoxicity in AD) neurons co-cultured with 11-12 mo 5xFAD astrocytes exhibited significantly higher MMP, compared to that with 5-6 mo mice. The 11-12 mo WT astrocytes were not observed to induce similar alterations in the neuronal MMP under vehicle or glutamate treatment. Prolonged high MMP has been reported to increased production of mitochondrial reactive oxygen species (ROS) and up-regulate autophagy, especially in neurons treated with glutamate (Kumari et al., 2012). Interestingly, both the WT and 5xFAD astrocytes isolated from 11-12 mo mice were incapable of buffering neuronal viability from the effects of glutamate exposure (Fig 4B). However, treating the co-cultures of neurons and astrocytes isolated from 5-6 mo mice WT or 5xFAD mice with glutamate had no effect on the viability of the neurons, suggesting an age-dependent reduction in astrocyte functions relating to neuronal support. Furthermore, the 11-12 mo 5xFAD astrocytes secreted reduced levels of anti-inflammatory interleukin 10 (IL-10) and interferon gamma (IFN-γ) in comparison to age-matched WT astrocytes (Supplementary Tables 2 and 3).

**Figure 4.**
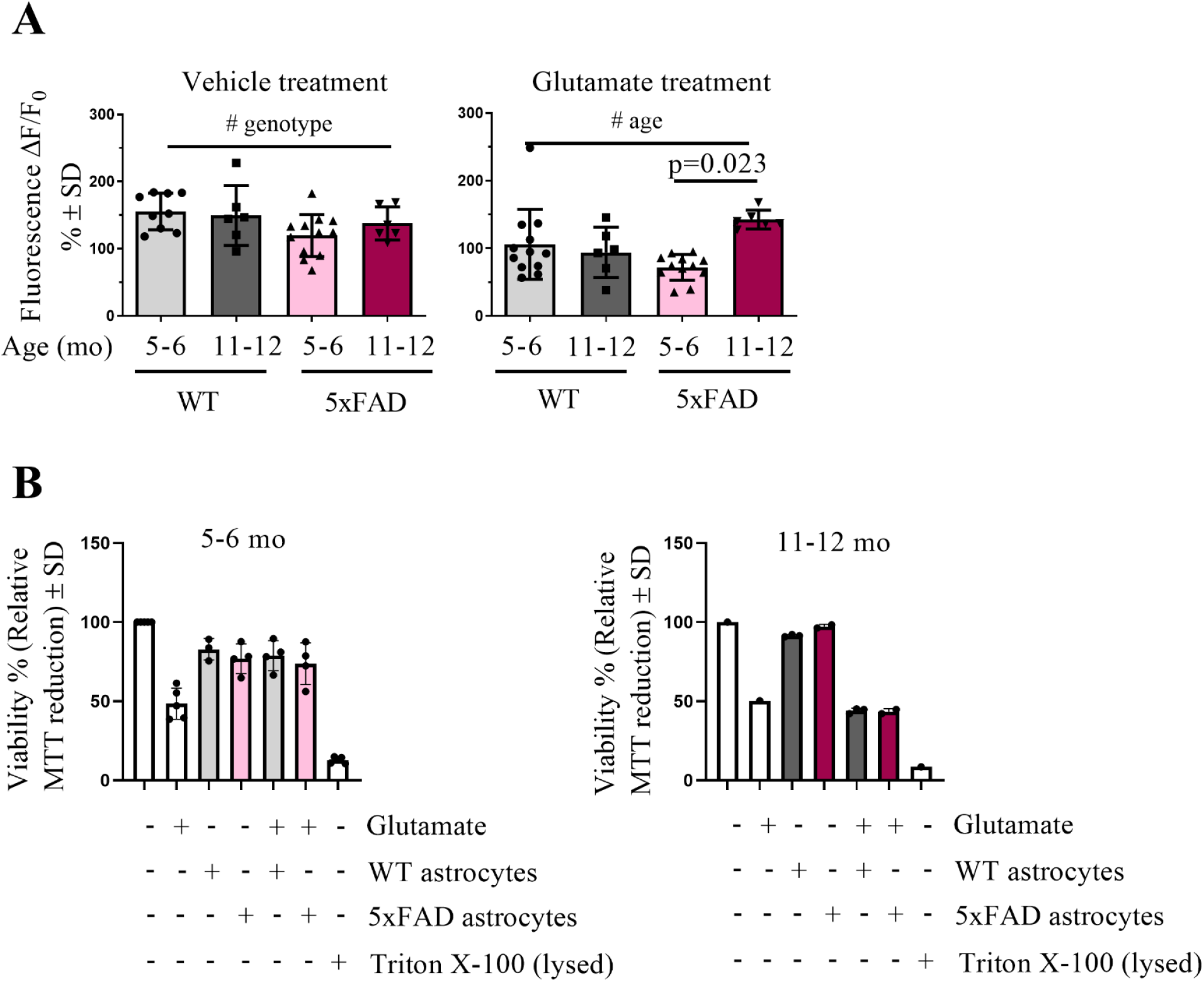
Astrocytes’ ability to support neuronal functions is altered in AD. (A) Mitochondrial membrane potential of primary WT neurons co-cultured with WT or 5xFAD adult astrocytes was measured by live cell imaging with rhodamine123 dye. The astrocytes were cultured on inserts on top of the neurons for 2 days prior the experiment and co-cultures were treated with 250 µM glutamate for 24 h before the live cell imaging. The bar graph shows the cells response to FCCP as ΔF/F_0_ (normalized to baseline), % ± SD. For neurons co-cultured with astrocytes harvested from 5-6 mo mice the data is shown as an average of 3-4 individual experiments with 3 technical replicates in each (n=3-4 mice per group in total). For astrocytes harvested from 10-12 mo mice, data shown as an average of 2 experiments with 3 technical replicates in each (N=2 mice per group in total). All graphs represent the mean ± SD. Two-way ANOVA genotype effect p=0.0495 for vehicle treated cultures and age-effect for the glutamate treated cultures p=0.0274. (B) The viability of the neurons was assessed with the MTT assay. The histograms show relative MTT reduction in neurons ± SD when neurons were co-cultured with or without astrocytes isolated from 5-6 mo or 11-12 mo) WT or 5xFAD mice for 2 days prior the experiment. For 24 h before the MTT assay all the cultures were treated with 250 µM glutamate or vehicle solution. Data is shown as an average of 3-5 individual experiments with 5-6 mo astrocytes and 1 experiment with 11-12 mo astrocytes (N=3-4 mice for astrocytes isolated from 5-6 mo mice, N= 2-3 mice for astrocytes isolated from 11-12 mo mice). One-way ANOVA.

### 3.5. Protein levels for mitophagy-inducer Ambra1 are significantly increased in aged 5xFAD astrocytes

To decipher whether autophagy-related processes are responsible for the observed transmitophagy alteration in aged AD astrocytes we assessed protein levels of common autophagy markers by Western blot. Immunoblotting against mitophagy- and autophagy-related proteins, p62 and LC3b, did not reveal significant changes in in astrocytes during aging or in AD (Fig. 5B). However, a two-fold increase in the protein level of Ambra1, a mitophagy inducer, was observed in aged (11-12 mo) 5xFAD astrocytes compared to WT astrocytes (Fig. 5A). The mRNA expression of *Ambra1* was not altered between the genotypes (data not shown).

**Figure 5.**
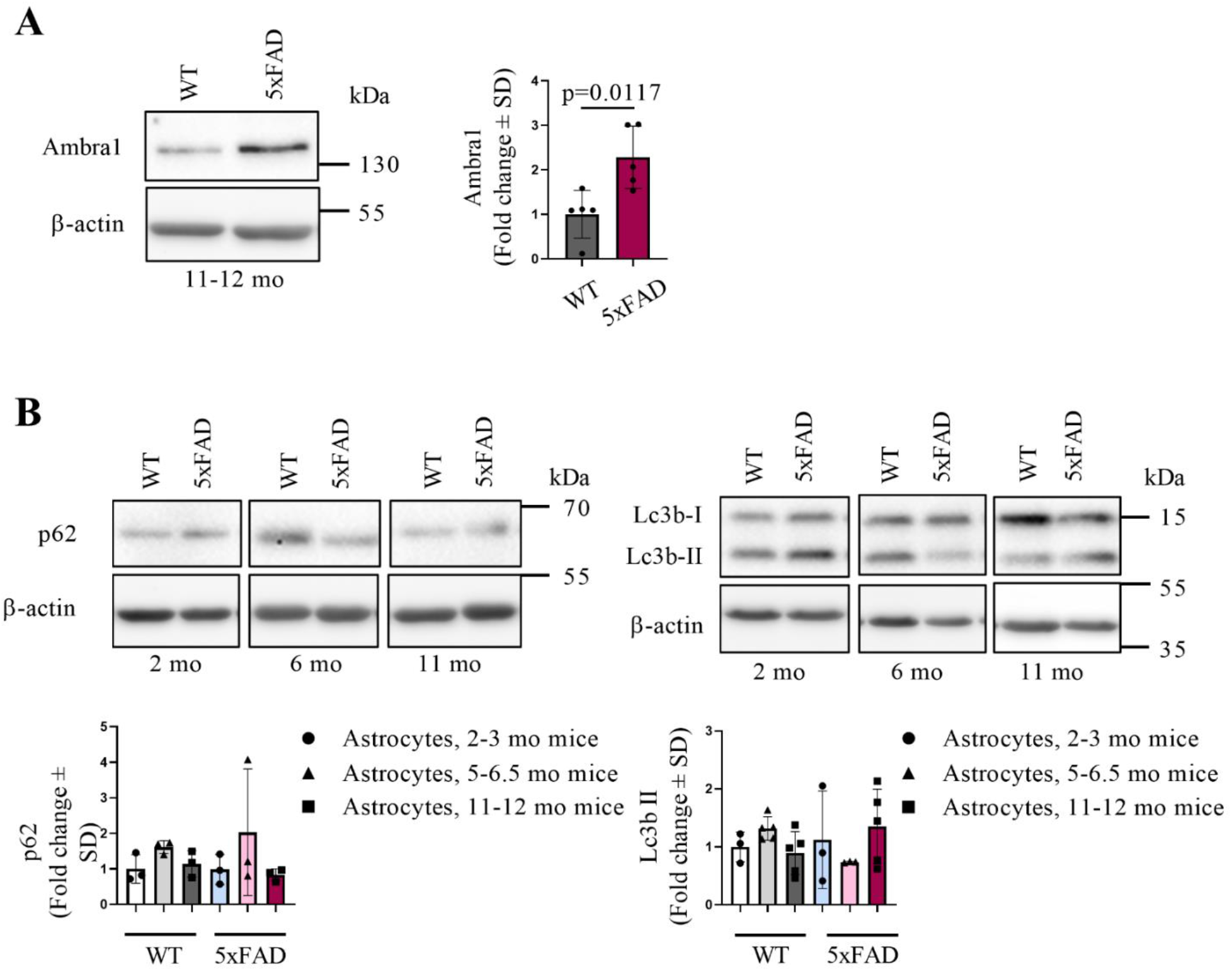
Expression of mitophagy-inducer Ambra1 is increased selectively in aged AD astrocytes. (A) The relative protein level of Ambra1 was assessed by Western blotting in adult astrocytes. N=3-5 mice per group. Unpaired two-tailed t test. (B) The relative protein levels of p62 and LC3B were assessed by Western blotting in adult astrocytes. N=3-5 mice per group. The ratio of protein expression was determined by normalizing the level of each protein to that of β-actin and the normalized value ratio to that of the value of 2-3 mo WT mice. In bar plots one dot represent astrocytes harvested from a single mouse.

## 4. Discussion

Many previously accepted facts about mitochondria have been recently re-evaluated, including the assumptions that each cell degrades its own mitochondria (Davis et al., 2014), and that mitochondria and their DNA are exclusively maternally inherited (Luo et al., 2018). These new findings have provided more on the importance of this organelle for previously unappreciated functions. The results presented herein demonstrate for the first time both in mouse and human that mitochondrial transfer occurs between neurons and AD-affected astrocytes, and that AD astrocytes degrade mitochondria derived from neurons. Moreover, mitochondrial transcellular movement and transmitophagy are impaired in AD.

Experiments with astrocytes treated with Aβ peptides have reported the induction of multiple alterations in astrocytic mitochondrial function (Abramov et al., 2004; Sarkar et al., 2014), but there are not many reports of mitochondrial functions of astrocytes isolated from mice modeling AD. Mitochondrial fractions of neonatal 5xFAD astrocytes have been shown to contain altered levels of metabolites and enzymatic activity related to the glycolytic pathway and TCA cycle. Similar changes were observed in WT astrocytes after exposure to oligomeric Aβ. (van Gijsel-Bonnello et al., 2017) One could think that defects in mitochondrial health of AD-affected astrocytes leads to increased internalization of neuronal mitochondria, for example due to an attempt to correct an energy deficit. However, our studies on the mitochondrial functions of astrocytes isolated from adult 5xFAD mouse brains did not reveal significant differences when compared to age-matched WT astrocytes in any of the assays we performed.

TNTs, thin elongations of the cell membrane consisting of F-actin, serve as highways for organelle transfer, including mitochondria, between cells (Rustom et al., 2004; Spees et al., 2006) Our data demonstrates the formation of TNT-like structures between neurons and astrocytes, and suggests they may mediate the transfer of mitochondria between these cell types. Although not assessed in the current paper, previous studies have shown that astrocytes from neonatal APP/PS1 mice form more TNTs than their WT counterparts (Sun et al., 2012) This finding supports our results and possibly explains, at least in part, our finding of increased mitochondrial transfer that occurs in aged 5xFAD astrocytes. The formation of TNTs between cells is driven by a concentration gradient of the protein S100a4, cleaved by caspase-3. The cell initiating the TNT formation has reduced levels of S100a4 compared to the recipient cells with higher S100a4 concentration (Sun et al., 2012). Interestingly, Yao et al. have observed increased caspase-3 activity in U87 astrocyte-like cells after treating the cells with Aβ (Yao et al., 2018).

The presence of AD pathology manifested as increased levels of brain Aβ_1-42_ could be speculated to activate caspase-3, leading to reduced levels of S100a4 in affected cells. Our results suggest that S100a4 levels are reduced in aged 5xFAD hippocampal neurons. On the other hand, astrocytes in the aged 5xFAD brain and 11-12 mo 5xFAD astrocytes cultured *in vitro* display increased S100a4. These findings imply that the aged 5xFAD astrocytes function as recipient cells for TNT-like structures formed by neurons, since the S100a4 protein gradient formed between these cell types would allow the transfer of mitochondria along TNT-like structures. This is in line with our observation of increased internalization of neuronal mitochondria by 5xFAD astrocytes over the WT astrocytes. Previously studies have also shown astrocytic S100a4 to be increased after head trauma or astrocytes under stress (Dmytriyeva et al., 2012; Kozlova and Lukanidin, 2002). The protein has been shown to be neuroprotective, acting partly via binding to the IL-10 receptor and via JAK/STAT3 pathway (Dmytriyeva et al., 2012). In addition, the authors suggested that S100a4 may be potent for activating the IL10R/Akt pathway. In other studies IL-6, IL-8 and especially IL-7 have been reported to function via JAK/STAT3 pathway and stimulate secretion of S100a4 from human chondrocytes (Yammani et al., 2009). In addition, when stimulated, adult 5xFAD astrocytes have been reported to secrete threefold more IL-6 compared to WT astrocytes (Iram et al., 2016). We did not observe differences in levels of secretion of IL-6 from unstimulated aged WT and 5xFAD astrocytes. Our result of increased S100a4 in astrocytes of the 12 mo 5xFAD mouse brain together with reduced secretion of anti-inflammatory IL-10 may, however, imply the involvement of the S100a4-IL-10 axis in AD astrocytes and in mitochondrial transfer between neurons and astrocytes.

Transmitophagy has been shown to occur in rodent astrocytes *in vivo* adjacent to the optic nerve head (Davis et al., 2014) and between dopaminergic neurons and their neighboring in the context of Parkinson’s disease (Morales et al., 2020). Our data are the first to demonstrate transmitophagy of neuron-derived mitochondria by human iPSC-derived astrocytes, and alterations to this process upon aging in AD astrocytes. Our data also supports the existence of transmitophagy both *in vitro* and *in vivo*. Altered transmitophagy was not associated with astrocytic phagocytosis capacity and basic mitochondrial functions, yet the cytochrome c oxidase activity appeared to be altered specifically around beta-amyloid plaques in aged 5xFAD brains and the aged AD astrocytes had significantly increased Ambra1 levels when compared to their WT counterparts. Existing literature supports the hypothesis of altered COX-activity in 5xFAD mice brains. Devi and Ohno (2012) have shown increased release of mitochondrial cytochrome c to cell cytosol in 5xFAD mice brains compared to the WT brains, regardless of the mouse age. In addition, the authors have described a trend (p=0.06) towards reduced COX activity in 12 month-old 5xFAD mouse brains (Devi and Ohno, 2012). Mitophagy induction via Ambra1 by its binding to LC3 is known to occur independently of the classical mitophagy-related protein Parkin (Strappazzon et al., 2015). Levels of p62 remained unaltered but the levels of LC3bII seemed to be slightly increased in the aged (11-12 mo) 5xFAD astrocytes. This suggests that unlike in synaptosomal (neuronal) mitochondria, in which aged 5xFAD cells display increased LC3bII and Parkin translocation (Wang et al., 2016), increased mitophagy in 5xFAD astrocytes is the result of up-regulated Ambra1 and its action together with LC3.

A key finding in this study is that a large amount of neuronal mitochondria are internalized and degraded by astrocytes in co-cultures. This finding agrees with a previous study reporting that degradation of large amounts of axonal mitochondria by glial cells in the optic nerve head (Davis et al., 2014). Notably, we also found mitochondria with both EGFP and mCherry signals inside the astrocytes, denoting mitochondria not yet present in an environment with a low pH suitable for degradation. Previous data indicated that the degrading neuronal mitochondria were found in astrocytes expressing high levels of marker for phagocytosis, Mac2 (Davis et al., 2014). On the other hand, reactive A1 type of astrocytes have been reported to be to present in AD-affected brains and during normal aging, showing reduced ability for neuronal support (Clarke et al., 2018; Liddelow et al., 2017). For example, in the article by Liddelow et al., astrocytes isolated from neonatal mice and induced to the A1 type by reactive microglia displayed reduced synapse engulfment and reduced gene expression of the phagocytosis related genes *Megf10* and *Mertk*. However, our data did not indicate alterations to phagocytosis by the aged 5xFAD astrocytes, suggesting that the phagocytosis process may not be responsible for the observed effects in transmitophagy and internalization of neuronal mitochondria.

Iram et al. have reported that astrocytes isolated from adult 5xFAD mouse brain show defects in neuronal support (Iram et al., 2016), and our results support this observation. Glutamate is known to strengthen the role of glycolysis as an energy source in cultured astrocytes (Yan et al., 2017). Our results indicated that glutamate treatment reduced neuronal viability when neurons were co-cultured with aged (11-12 mo) astrocytes despite the genotype of the astrocytes. However, the aged 5xFAD astrocytes failed to prevent the effects of glutamate excitotoxicity on neuronal MMP. The inability of neuronal mitochondria to reduce MMP upon glutamate exposure in a neuron-astrocyte co-culture indicates a dysfunctional response in the aged astrocytes, resulting in increased stress and loss of quality, thus direction for degradation including transmitophagy. Recently it was reported that only those retinal ganglion cells that were already damaged by a previous insult were prone to cellular death induced by neurotoxic astrocytes (Guttenplan et al., 2020). This is in line with our observations of the effects of aged and especially aged 5xFAD astrocytes on the neuronal health of glutamate treated neurons.

In the past, researchers have heavily relied on conducting experiments with astrocytes isolated from neonatal mice. In this study we utilized astrocytes isolated from the brains of adult mice at various age points and report significant age- and genotype-related differences in the performed assays, emphasizing the importance of using cells derived from aged animals especially when studying age-related diseases. Transmitophagy was also observed in human iPSC-derived astrocytes and *in vivo* in the mouse hippocampi. Further studies should focus on developing means to study transmitophagy *in vivo* in the human brain and to assess whether the observed alterations represent an early indication of AD-related pathology. Taken together, our results along with the existing data highlights the importance of mitochondrial traffic between neurons and astrocytes and demonstrates AD-induced alterations to astrocytic functions.

## Acknowledgements

We thank Mrs. Mirka Tikkanen, Ms. Laila Kaskela and Ms. Anna Palmgren for their expert technical assistance. The study was supported by Academy of Finland, Sigrid Juselius Foundation and University of Eastern Finland, and carried out with the support of UEF Cell and Tissue Imaging Unit, University of Eastern Finland, Finland. The authors thank Biocenter Kuopio Viral Gene Transfer service for providing lentiviral vectors for GFP.

## Disclosure

The authors have no actual or potential conflicts of interest.

## Data availability statement

Data available on reasonable request from the corresponding author.

## Notes

### Competing Interest Statement

The authors have declared no competing interest.

## References

Abramov, A. Y., Canevari, L., & Duchen, M. R. β-Amyloid Peptides Induce Mitochondrial Dysfunction and Oxidative Stress in Astrocytes and Death of Neurons through Activation of NADPH Oxidase. J Neurosci. 2004; 24: 565–75.

Clarke, L. E., Liddelow, S. A., Chakraborty, C., Münch, A. E., Heiman, M., & Barres, B. A. Normal aging induces A1-like astrocyte reactivity. Proc Natl Acad Sci U S A 2018; 115: E1896–E05.

Davis, C. H. O., Kim, K. Y., Bushong, E. A., Mills, E. A., Boassa, D., Shih, T., … Marsh-Armstrong, N. Transcellular degradation of axonal mitochondria. Proc Natl Acad Sci U S A 2014; 111: 9633–38.

Devi, L., Ohno M. Mitochondrial dysfunction and accumulation of the β-secretase-cleaved C-terminal fragment of APP in Alzheimer’s disease transgenic mice. Neurobiol Dis. 2012; 45:417–24.

Dmytriyeva, O., Pankratova, S., Owczarek, S., Sonn, K., Soroka, V., Ridley, C. M., Marsolais A., Lopez-Hoyos M., Ambartsumian N., Lukanidin E., Bock E., Berezin V., Kiryushko, D. The metastasis-promoting S100A4 protein confers neuroprotection in brain injury. Nat Commun. 2012; 3: 1197.

English, K., Shepherd, A., Uzor, N. E., Trinh, R., Kavelaars, A., Heijnen, C. J. Astrocytes rescue neuronal health after cisplatin treatment through mitochondrial transfer. Acta Neuropathol Commun. 2020; 8: 36.

Fang, E. F., Hou, Y., Palikaras, K., Adriaanse, B. A., Kerr, J. S., Yang, B., Lautrup S., M.M. Hasan-Olive Caponio D., Dan X., Rocktäschel P., Croteau D.L., Akbari M., Greig N.H., Fladby T., Nilsen H., Cader M.Z., Mattson M.P., Tavernarakis N., Bohr, V. A. Mitophagy inhibits amyloid-β and tau pathology and reverses cognitive deficits in models of Alzheimer’s disease. Nat Neurosci. 2019; 22: 401–12.

Foti, S. C., Hargreaves, I., Carrington, S., Kiely, A. P., Houlden, H., Holton, J. L. Cerebral mitochondrial electron transport chain dysfunction in multiple system atrophy and Parkinson’s disease. Sci Rep. 2019, 9, 6559.

Guttenplan, K. A., Stafford, B. K., El-Danaf, R. N., Adler D.I., Münch A.E., Weigel, M. K., Huberman, A. D., Liddelow, S. A. Neurotoxic Reactive Astrocytes Drive Neuronal Death after Retinal Injury. Cell Rep. 2020; 31: 107776.

Hayakawa, K., Esposito, E., Wang, X., Terasaki, Y., Liu, Y., Xing, C., Ji X., Lo, E. H. Transfer of mitochondria from astrocytes to neurons after stroke. Nature 2016; 535: 551–55.

Hong, Y., Liu, Y., Zhang, G., Wu, H., Hou, Y. Progesterone suppresses Aβ42-induced neuroinflammation by enhancing autophagy in astrocytes. Int Immunopharmacol. 2018; 54: 336–43.

Iram, T., Trudler, D., Kain, D., Kanner, S., Galron, R., Vassar, R., Barzilai A., Blinder P., Fishelson Z., Frenkel, D. Astrocytes from old Alzheimer’s disease mice are impaired in Aβ uptake and in neuroprotection. Neurobiol Dis. 2016; 96: 84–94.

Kanninen, K., Heikkinen, R., Malm, T., Rolova, T., Kuhmonen, S., Leinonen, H., Ylä-Herttuala S., Tanila H., Levonen A-L., Koistinaho M., Koistinaho, J. Intrahippocampal injection of a lentiviral vector expressing Nrf2 improves spatial learning in a mouse model of Alzheimer’s disease. Proc Natl Acad Sci U S A 2019; 106: 16505–10.

Konttinen H., Gureviciene I., Oksanen M., Grubman A., Loppi S., Huuskonen M.T., Korhonen P., Lampinen R., Keuters M., Belaya I., Tanila H., Kanninen K.M., Goldsteins G., Landreth G., Koistinaho J., Malm T. PPARβ/δ-agonist GW0742 ameliorates dysfunction in fatty acid oxidation in PSEN1ΔE9 astrocytes. Glia, 2019;67:146–59.

Kozlova, E. N., Lukanidin, E. Mts1 protein expression in the central nervous system after injury. Glia 2002; 37: 337–48.

Kumari, S., Mehta, S. L., Li, P. A. Glutamate induces mitochondrial dynamic imbalance and autophagy activation: Preventive effects of selenium. PLoS ONE 2012; 7: e39382.

Lampinen, R., Belaya, I., Boccuni, I., Malm, T., Kanninen, K. M. Mitochondrial Function in Alzheimer’s Disease: Focus on Astrocytes. In Astrocyte - Physiology and Pathology. Gentile, MT., editor. Intech open 2018; p. 139–162.

Lane, C. A., Hardy, J., Schott, J. M. Alzheimer’s disease. Eur J Neurol. 2018; 25: 59–70.

Liddelow S.A., Guttenplan K.A., Clarke L.E., Bennett F.C., Bohlen C.J., Schirmer L., Bennett M.L., Münch A.E., Chung W.S., Peterson T.C., Wilton D.K., Frouin A., Napier B.A., Panicker N., Kumar M., Buckwalter M.S., Rowitch D.H., Dawson V.L., Dawson T.M., Stevens B., Barres B.A. Neurotoxic reactive astrocytes are induced by activated microglia. Nature 2017; 541: 481–87.

Loppi S., Korhonen P., Bouvy-Liivrand M., Caligola S., Turunen T.A., Turunen M.P., Hernandez de Sande A., Kołosowska N., Scoyni F., Rosell A., García-Berrocoso T., Lemarchant S., Dhungana H., Montaner J., Koistinaho J., Kanninen K.M., Kaikkonen M.U., Giugno R., Heinäniemi M., Malm T. Peripheral inflammation preceeding ischemia impairs neuronal survival through mechanisms involving miR-127 in aged animals. Aging Cell. 2021;20:e13287.

Luo S, Valencia C.A, Zhang J, Lee N.C, Slone J, Gui B, Wang X, Li Z, Dell S, Brown J, Chen S.M, Chien Y.H, Hwu W.L, Fan P.C, Wong L.J, Atwal P.S, Huang T. Biparental inheritance of mitochondrial DNA in humans. Proc Natl Acad Sci U S A 2018, 115, 13039–44.

Morales, I., Sanchez, A., Puertas-Avendaño, R., Rodriguez-Sabate, C., Perez-Barreto, A., Rodriguez, M. Neuroglial transmitophagy and Parkinson’s disease. Glia 2020; 68, 2277–99.

Nachlas, M. M., Tsou, K. C., de Souza, E., Cheng, C. S., Seligman, A. M. Cytochemical demonstration of succinic dehydrogenase by the use of a new p-nitrophenyl substituted ditetrazole. J Histochem Cytochem. 1957; 5: 420–36.

Oakley, H., Cole, S. L., Logan, S., Maus, E., Shao, P., Craft, J., Guillozet-Bongaarts A., Ohno M., Disterhoft J., Van Eldik L., Berry, R., Vassar, R. Intraneuronal β-amyloid aggregates, neurodegeneration, and neuron loss in transgenic mice with five familial Alzheimer’s disease mutations: Potential factors in amyloid plaque formation. J Neurosci. 2006; 26: 10129–40.

Oksanen, M., Petersen, A. J., Naumenko, N., Puttonen, K., Lehtonen, Š., Gubert Olivé, M., Shakirzyanova A., Leskelä S., Sarajärvi T., Viitanen M., Rinne J.O., Hiltunen M., Haapasalo A., Giniatullin R., Tavi P., Zhang S., Kanninen K.M., Hämäläinen R.H., Koistinaho, J. PSEN1 Mutant iPSC-Derived Model Reveals Severe Astrocyte Pathology in Alzheimer’s Disease. Stem Cell Reports 2017; 9: 1885–97.

Pickett, E. K., Rose, J., McCrory, C., McKenzie, C. A., King, D., Smith, C., Gillingwater T.H., Henstridge C.M., Spires-Jones, T. L. Region-specific depletion of synaptic mitochondria in the brains of patients with Alzheimer’s disease. Acta Neuropathol. 2018; 136: 747–57.

Pickford, F., Masliah, E., Britschgi, M., Lucin, K., Narasimhan, R., Jaeger, P. A., Small S., Spencer B., Rockenstein E., Levine B., Wyss-Coray, T. The autophagy-related protein beclin 1 shows reduced expression in early Alzheimer disease and regulates amyloid β accumulation in mice. J Clin Invest. 2008; 118: 2190–99.

Reddy, P. H., Oliver, D. M. Amyloid Beta and Phosphorylated Tau-Induced Defective Autophagy and Mitophagy in Alzheimer’s Disease. Cells 2019; 8: 488.

Rustom, A., Saffrich, R., Markovic, I., Walther, P., Gerdes, H. H. Nanotubular Highways for Intercellular Organelle Transport. Science 2004; 303: 1007–10.

Sarkar, P., Zaja, I., Bienengraeber, M., Rarick, K. R., Terashvili, M., Canfield, S., Falck J.R., Harder, D. R. Epoxyeicosatrienoic acids pretreatment improves amyloid-induced mitochondrial dysfunction in cultured rat hippocampal astrocytes. Am J Physiol Heart Circ Physiol 2014; 306: 475–84.

Soundara Rajan, T., Gugliandolo, A., Bramanti, P., Mazzon, E. Tunneling Nanotubes-Mediated Protection of Mesenchymal Stem Cells: An Update from Preclinical Studies. Int J Mol Sci. 2020; 21: 3481.

Spees, J. L., Olson, S. D., Whitney, M. J., Prockop, D. J. Mitochondrial transfer between cells can rescue aerobic respiration. Proc Natl Acad Sci U S A 2006; 103: 1283–88.

Strappazzon, F., Nazio, F., Corrado, M., Cianfanelli, V., Romagnoli, A., Fimia, G. M., Campello S., Nardacci R., Piacentini M., Campanella M., Cecconi, F. AMBRA1 is able to induce mitophagy via LC3 binding, regardless of PARKIN and p62/SQSTM1. Cell Death Differ. 2015; 22: 419–32.

Sun, X., Wang, Y., Zhang, J., Tu, J., Wang, X. J., Su, X. D., Wang L., Zhang, Y. Tunneling-nanotube direction determination in neurons and astrocytes. Cell Death Dis.2012; 3: e438– e38.

Tiihonen, J., Koskuvi, M., Lähteenvuo, M., Virtanen, P. L. J., Ojansuu, I., Vaurio, O., Gao Y., Hyötyläinen I., Puttonen K.A., Repo-Tiihonen E., Paunio T., Rautiainen M.R., Tyni S., Koistinaho J., Lehtonen, Š. Neurobiological roots of psychopathy. Mol Psychiatry. 2019;25: 3432–41

van Gijsel-Bonnello, M., Baranger, K., Benech, P., Rivera, S., Khrestchatisky, M., de Reggi, M., Gharib, B. Metabolic changes and inflammation in cultured astrocytes from the 5xFAD mouse model of Alzheimer’s disease: Alleviation by pantethine. PLoS ONE 2017; 12: e0175369.

Wang, L., Guo, L., Lu, L., Sun, H., Shao, M., Beck, S. J., Li L., Ramachandran J., Du Y., Du, H. Synaptosomal mitochondrial dysfunction in 5xFAD mouse model of Alzheimer’s disease. PLoS ONE 2016; 11: e0150441.

Wang, W., Zhao, F., Ma, X., Perry, G., & Zhu, X. Mitochondria dysfunction in the pathogenesis of Alzheimer’s disease: recent advances. Mol Neurodegener. 2020; 15: 30.

Yammani, R. R., Long, D., Loeser, R. F. Interleukin-7 stimulates secretion of S100A4 by activating the JAK/STAT signaling pathway in human articular chondrocytes. Arthritis Rheum. 2009; 60: 792–800.

Yan, X., Shi, Z. F., Xu, L. X., Li, J. X., Wu, M., Wang, X. X., Jia M., Dong L.P. Yang S.H., Yuan, F. Glutamate Impairs Mitochondria Aerobic Respiration Capacity and Enhances Glycolysis in Cultured Rat Astrocytes. Biomed Environ Sci. 2017; 30, 44–51.

Yao, Y., Huang, J. Z., Chen, Y., Hu, H. J., Tang, X., Li, X. Effects and mechanism of amyloid β1-42 on mitochondria in astrocytes. Mol Med Rep. 2018; 17: 6997–74.

Ye, X., Sun, X., Starovoytov, V., Cai, Q. Parkin-mediated mitophagy in mutant hAPP neurons and Alzheimer’s disease patient brains. Hum Mol Genet. 2015; 24: 2938–51.

