## Supplementary material and methods for "Alzheimer’s disease alters astrocytic functions related to neuronal support and transcellular internalization of mitochondria"

### Cytochrome c oxidase activity

For assessing the cytochrome c oxidase (COX) activity in astrocytes surrounding the beta-amyloid plaques, the 12 mo mice were deeply anesthetized and perfused transcardially with heparinized saline. The brain hemispheres were stored at -70℃ before cryosectioning to 10 µm sections with a 150 µm interval. The COX/SDH staining was performed as previously described (Foti et al., 2019; Nachlas, Tsou, de Souza, Cheng, & Seligman, 1957) with minor modifications. Briefly, the sections were rehydrated with 0.05M Na phosphate buffer (pH 7.4) before incubating them in substrate buffer for the COX assay (0.05 M Na phosphate buffer (pH 7.4) containing 1 mg/ml cytochrome C (Sigma-Aldrich), 75 mg/ml sucrose, 1 mg/ml DAB (Sigma-Aldrich), 0.8 mg/ml catalase (Sigma-Aldrich) for 1h at 37℃. After incubation the sections were washed with $dH_2O$. For the succinic dehydrogenase (SDH) assay, the sections were incubated directly after COX incubating medium in SDH staining solution. Briefly, equal volumes of 0.2 M Na phosphate buffer (pH 7.6) and 0.2 M sodium succinate (Sigma-Aldrich) were combined. Next, when starting the incubation, the buffered substrate reagent was combined 1:1 with aqueous solution containing 1mg/ml nitrotetrazolium blue (Sigma-Aldrich). The sections were incubated in SDH staining solution for 1.75h at 37℃. After SDH staining the sections were washed first once with 0.05M PB and then once more with PBS containing 0.2% Tween20. Next, the sections were blocked with mouse on mouse blocking reagent (Vector Laboratories) for 40 minutes at room temperature, following washes with PBS containing 0.2% Tween20 and blocking with 10% normal goat serum prepared in PBS containing 0.2% Tween20 for 1h at room temperature before incubation overnight at room temperature with the mixture of anti-GFAP (Dako Z033429-2, 1:500) and anti-amyloid β, clone WO-2 (Millipore, Burlington, MA, USA, MABN10, 1:1000) primary antibodies diluted in PBST. Next, the sections were washed with PBS containing 0.2% Tween20 and incubated with Alexa Fluor 488 and Alexa Fluor 568 (both 1:300, Thermo Fisher Scientific) secondary antibodies diluted in 5% normal goat serum in PBS containing 0.2% Tween20 for 2 h at room temperature. Finally, the sections were embedded with Vectashield mounting medium (Vector Laboratories) for fluorescence with 4′,6-diamidino-2-phenylindole (DAPI). Stained sections were imaged with Zeiss Axio Imager 2 fluorescent microscope with 20x and 40x objective.

Supplementary file, Material and methods for supplementary files

**Analysis of mitochondrial metabolism**

The MitoStress Test was performed according to the manufacturer's instructions (Agilent, Santa Clara, CA, USA). The astrocytes were seeded on cell culture microplates as 15000 cells/XFe96-well two days before the assay. The XF base assay medium was supplemented with 25 mM D-glucose (Sigma-Aldrich), 2 mM sodium pyruvate (Gibco), 2 mM GlutaMAX (Thermo Fisher Scientific) and the pH of the medium was adjusted to 7.4. The oxygen consumption rates (OCR) following injections with oligomycin, FCCP, rotenone and antimycin A (all at a final concentration of 1 µM, Sigma-Aldrich) were recorded with Seahorse XFe96 Analyzer (Agilent). The total protein concentrations/ well were measured with Pierce BCA protein assay kit (Thermo Fisher Scientific) from cells lysed in 1xRIPA buffer and the data was analyzed utilizing Wave 2.6.0 (Agilent).

**Quantification of intracellular ATP levels**

Astrocytes were seeded as 40000 cells/48-well for one day prior the assay. The intracellular levels of ATP in astrocytes were analyzed using the ATPLite Luminescence Assay System (Perkin Elmer) according to the manufacturer's instructions. Luminescence was read with the Wallac Victor 1420 microplate reader (Perkin Elmer).

**Phagocytosis assay**

One vial of pHrodoTM Green Zymosan BioParticlesTM Conjugate (Thermo Fisher Scientific) was resuspended in 4 ml of medium (Opti-Mem or DMEM/F-12, GlutaMAX™ Supplement) and briefly vortexed and sonicated for 5 minutes. The cell culture medium was removed from astrocytes seeded one day prior the assay on 96-wells as 10000 cells/well and quickly replaced with 100 µl of the prepared bioparticle suspension. As a control, one well in each condition received medium without bioparticles. Cells were incubated at 37℃ in 5% CO2 for 2 hours. After the incubation, the solution was removed, and cells were washed twice with the cell culture medium. Bisbenzimide (Sigma-Aldrich, 1:2000 or 1:3500 diluted in cell culture medium) was added for 5 minutes for staining the nuclei and then replaced with fresh culture medium.

Supplementary file, Material and methods for supplementary files

Live imaging was performed on the Zeiss AxioObserver Z1 fluorescent microscope and with Zen 2 (blue edition) software. 4-5 images per well were taken from all 5 replicates and controls using a 10x objective. Images were analyzed with ImageJ. Phagocytosis was quantified as follows area of beads (%) / area of nuclei (%) *100. Each experiment was carried out three times.

**Cytometric bead array**

For assessing the secreted levels of IL-6, IL-10, MCP-1, IFN- γ, TNF and IL-12p70 from the adult mouse astrocytes isolated from 5-6 mo and 11-12 mo WT or 5xFAD mice, the astrocytes were plated on 48-well plates at 12500 cells/well and grown for five days before collecting the cell culture supernatant for cytometric bead array (CBA). The CBA assay (CBA Mouse Inflammation Kit #552364, BD Biosciences, San Jose, CA, USA) was performed according to the manufacturer's instructions using CytoFLEX S flow cytometer (Beckman Coulter, Indianapolis, IN, USA) for data acquisition. Data was analyzed with FCAP Array™ v2.0.2 151 software (SoftFlow Hungary Ltd, Pecs, Hungary).

**Quantitative RT-PCR**

For collection of total RNA, the astrocytes were seeded on 6-well plates at 100000 - 150000 cells/well for three days prior RNA extraction. Cells were washed with DPBS (Thermo Fisher Scientific) and the RNA was isolated using TRI reagent (Sigma-Aldrich) according to the manufacturer's protocol. 0.5 µg of RNA was reverse transcribed with High Capacity Reverse Transcription kit or with Maxima Reverse Transcriptase (all ThermoFisher Scientific) according to manufacturer's instructions.

For relative expression of the mRNA encoding for selected mouse genes were quantified according to the manufacturer's instructions for Applied Biosystems™ StepOne™ Real-Time PCR System using TaqMan® real-time PCR assay mixes (Thermo Fisher Scientific) for targets (Supplementary table 1). The expression levels of these genes were normalized to S18 ribosomal RNA and presented as log2 fold change to the age-matched WT mice.

Supplementary file, Material and methods for supplementary files

**Enzyme-linked immunosorbent assay**

Cell lysates for ELISA were collected from 5-6 mo and 11-12 mo adult astrocytes from the same wells as the media samples for cytometric bead array (CBA). The cells were lysed in 1x RIPA buffer supplemented with 1x cOmplete protease inhibitor cocktail (Roche, Basel, Switzerland) and analysed with Mouse Protein S100-A4(S100A4) ELISA kit (Cusabio, Houston, TX, USA #CSB-EL020632MO) as instructed in the kit's manual. Total amounts of protein in cell lysate samples were quantified with BCA assay (Pierce™ BCA Protein Assay Kit, Thermo Scientific) according to the manufacturer's instructions. The results were calculated as pg/mg protein ± SD.
