## Supplementary figures and tables for "Alzheimer’s disease alters astrocytic functions related to neuronal support and transcellular internalization of mitochondria"

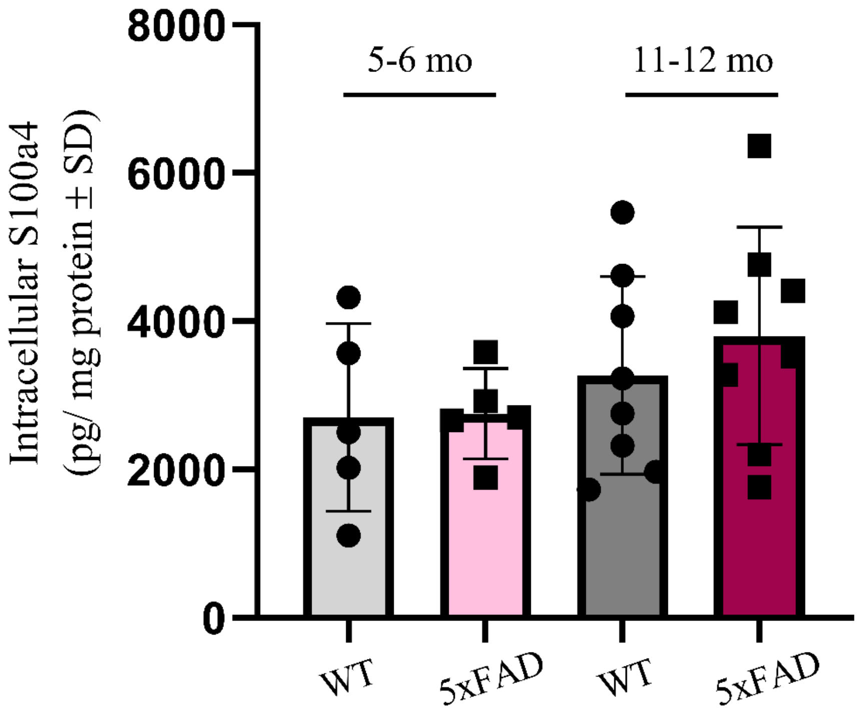


**Supplementary Figure 1. Levels of intracellular S100a4 seem to be highest in astrocytes harvested from 11-12 mo 5xFAD mice compared to WT astrocytes or astrocytes derived from 5-6 mo mice.** The S100a4 levels were assessed in adult astrocyte samples with ELISA. Each dot represents one single mouse where the astrocytes were harvested. Two-way ANOVA * $P \leq 0.05$, * age (p=0.0283).

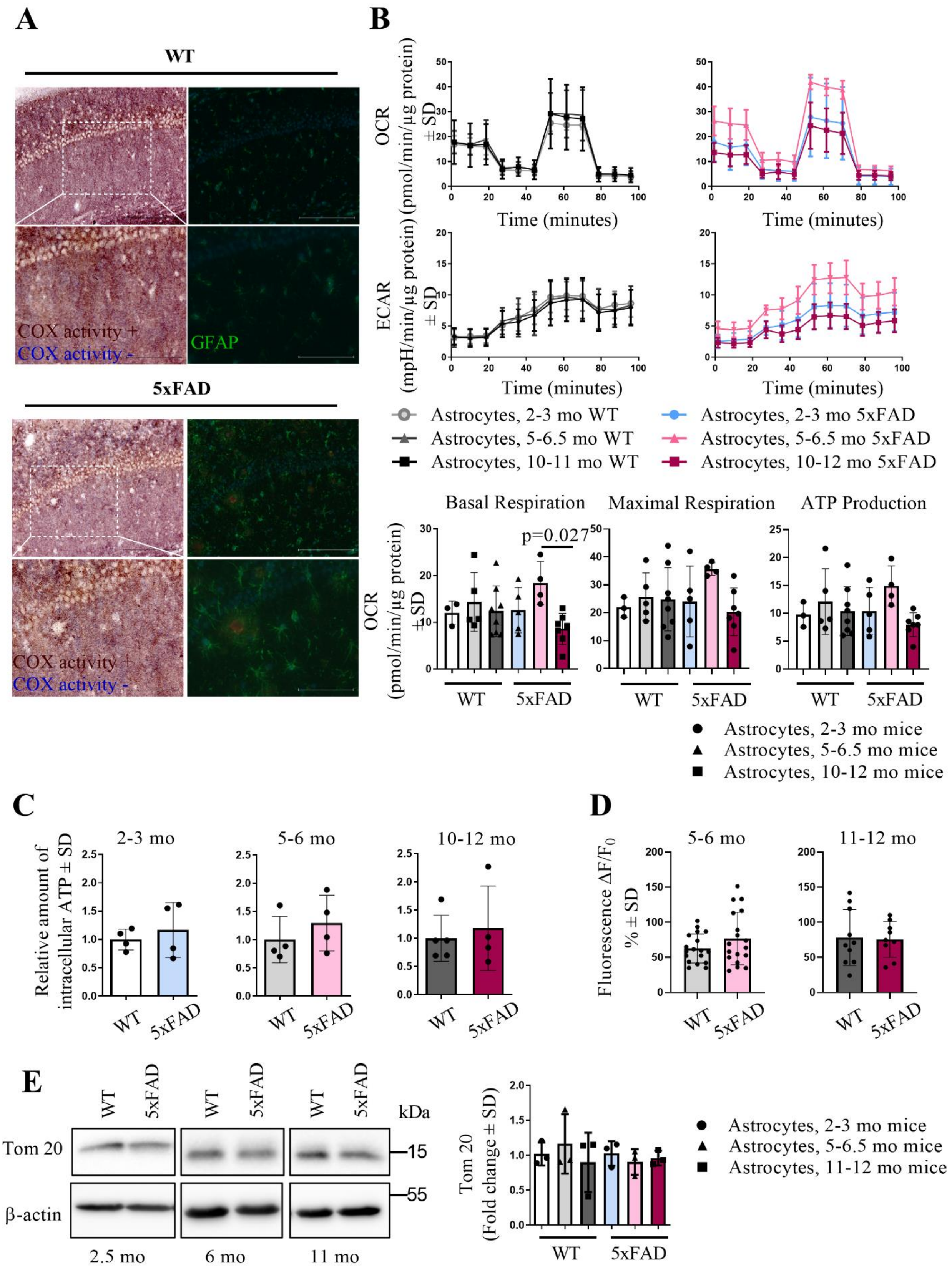


**Supplementary Figure 2. Mitochondrial functions of astrocyte's own mitochondria are**

**unaltered in aging and in AD** (A) COX/SDH immunohistochemical staining combined with anti-GFAP and anti-β-amyloid staining in sagittal sections of 12 mo WT and 5xFAD mouse hippocampus. Example images with 20x objective (scale bar 200 µm) and boxed areas with 40x objective (scale bar 100 µm). The cells with impaired cytochrome c activity are visualized in blue, indicating succinate dehydrogenase (SDH) activity without cytochrome c activity, and co-localized with astrocytes immunostained with an anti-GFAP antibody in green. The brown cells indicate those with normal activity of cytochrome c and areas immunostained with anti-β-amyloid antibody are shown in red. (B) Mitochondrial respiration rates of adult astrocytes assessed with Seahorse MitoStress test (Agilent) astrocytes harvested from WT and 5xFAD brains. Two-way ANOVA. Unpaired two-tailed t test was used for OCR values between 5-6.5 mo and 10-12 mo 5xFAD astrocytes. (C) Intracellular ATP levels in adult astrocytes harvested from the 5xFAD mice when compared to adult astrocytes isolated from the age-matched WT mice. Unpaired two-tailed t test. (D) Mitochondrial membrane potential of astrocytes harvested from 5-6 mo and 10-11 mo WT and 5xFAD mouse brains. Unpaired two-tailed t test. (E) Immunoblotting with anti-Tom20 in adult astrocytes. Two-way ANOVA. (A) Example images; (B) Graph shows the average of 5 individual MitoStress test assays with N=3-8 mice/group in total. In bar plots one dot represents astrocytes derived from one mouse. (C) Average of results from 4-5 biologically individual experiments with 3-6 technical replicates in each, data points represent the average result for astrocytes isolated from each individual mice in these experiments; (D) For astrocytes harvested from 5-6 mo mice the data is shown as an average of 6 individual experiments with 3 technical replicates in each (n=6 mice per group in total). For astrocytes harvested from 10-11 mo mice, data are shown as an average of 2 mice with 4-5 technical replicates in each (N=2 mice per group in total); (E) N=3 mice per group. In the bar plot one dot represents astrocytes harvested from a single mouse. All graphs represent the mean ± SD.

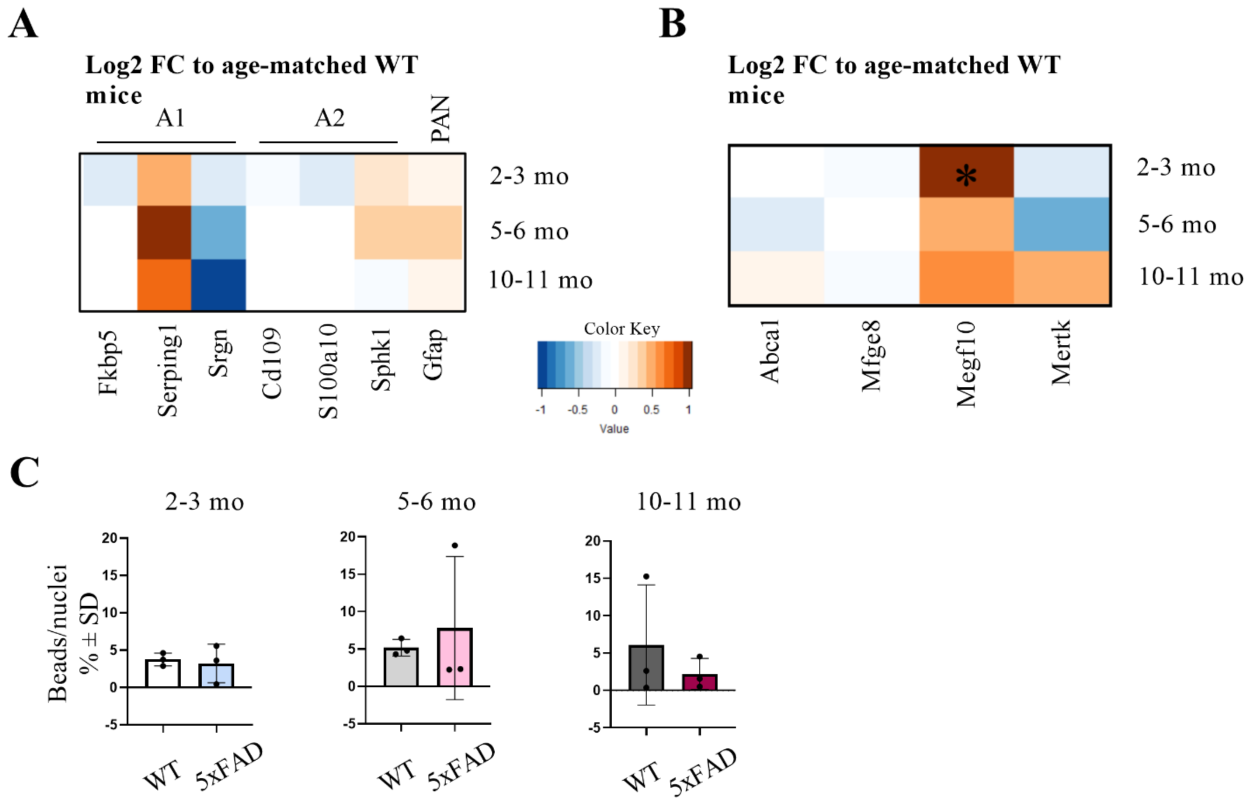


**Supplementary Figure 3. Astrocyte reactivity and phagocytosis are unchanged upon aging or in AD** (A) Heatmap of WT and 5xFAD adult astrocyte activity state assessed with RT-qPCR. The ratio of mRNA expression for genes of interest was determined by normalizing to the level of mRNA to that of eukaryotic ribosomal protein S18 and the normalized values of 5xFAD astrocytes as log2 fold changes to age-matched WT mice. N=3-7 mice per group with 3-6 technical replicates per mouse. Unpaired two-tailed t test. (B) Heatmap of expression of phagocytosis-related genes assessed with RT-qPCR. The ratio of mRNA expression for genes of interest was determined by normalizing to the level of mRNA to that of eukaryotic ribosomal protein S18 and the normalized values of 5xFAD astrocytes as log2 fold changes to age-matched WT mice. Asterisk marks significant difference for Megf10 (Log2 FC 1.0689, $p<0.045$) between WT and 5xFAD astrocytes isolated from 2-3 mo mice. N=3-7 mice per group with 2-6 technical replicates per mouse. Unpaired two-tailed t test. (C) Phagocytic capacity of astrocytes was measured by using of pHrodo Green Zymosan Bioparticles Conjugates, which were added to cells for 2 h. The histograms show the percentage of the ratio area of beads/ area of nuclei ± SD. The average of 3 biologically individual experiments, each data point represents astrocytes derived from a single mouse. Unpaired two-tailed t test.

**Supplementary table 1. Primers used for RT-qPCR**

| Gene | Supplier | Product catalog number and ID |
|---|---|---|
| Abca1 | Thermo Fisher Scientific | 4331182, Mm00442646_m1 |
| Ambra1 | Thermo Fisher Scientific | 4331182, Mm00554370_m1 |
| Cd109 | Thermo Fisher Scientific | 4331182, Mm00462151_m1 |
| Fkbp5 | Thermo Fisher Scientific | 4331182, Mm00487406_m1 |
| Gfap | Thermo Fisher Scientific | 4331182, Mm01253033_m1 |
| Hmox1 | Thermo Fisher Scientific | 4331182, Mm00516005_m1 |
| Mfge8 | Thermo Fisher Scientific | 4331182, Mm00500549_m1 |
| Megf10 | Thermo Fisher Scientific | 4331182, Mm01257625_m1 |
| Mertk | Thermo Fisher Scientific | 4331182, Mm00434920_m1 |
| Nqo1 | Thermo Fisher Scientific | 4331182, Mm01253561_m1 |
| S100a4 | Thermo Fisher Scientific | 4331182, Mm00803371_m1 |
| S100a10 | Thermo Fisher Scientific | 4331182, Mm00501457_m1 |
| Serping1 | Thermo Fisher Scientific | 4331182, Mm00437835_m1 |
| Sphk1 | Thermo Fisher Scientific | 4331182, Mm00448841_g1 |
| Srgn | Thermo Fisher Scientific | 4331182, Mm01169070_m1 |
| Eukaryotic 18S rRNA | Thermo Fisher Scientific | 4333760F, Hs99999901_s1 |

**Supplementary Table 2. Effect of AD on cytokine secretion from astrocytes isolated from 5-6 mo WT and 5xFAD mice brain.** Cytokines secreted by astrocytes isolated from 5-6 mo WT and 5xFAD mouse brains. Data are shown as mean ± SD. Unpaired t-test. N=5 mice per group with 4 technical replicates, except WT for IL-6 and 5xFAD for MCP-1 N=4 mice.

| Cytokine | Cytokine concentration (pg/ml) | | Unpaired t-test | |
|---|---|---|---|---|
| | WT 5-6 mo (N=5) | 5xFAD 5-6 mo (N=5) | p-value | Significanc e |
| IL-6 | 4.0 ± 5.1 | 8.6 ± 8.5 | 0,4345 | ns |
| IL-10 | 47.6 ± 24.2 | 34.0 ± 30.1 | 0,5011 | ns |
| MCP-1 | 0.1 ± 0.1 | 0.02 ± 0.04 | 0,3054 | ns |
| IFN-γ | 3447.4 ± 1167.6 | 2382.6 ± 1394.9 | 0,2754 | ns |
| TNF | 2.3 ± 0.9 | 1.9 ± 1.3 | 0,6962 | ns |
| IL-12p70 | 2249.6 ± 1084.1 | 1481.3 ± 819.1 | 0,2909 | ns |

**Supplementary Table 3. Effect of AD on cytokine secretion from astrocytes isolated from 11-12 mo WT and 5xFAD mice brain.** Cytokines secreted by astrocytes isolated from 11-12 mo WT and 5xFAD mouse brains. Data are shown as mean ± SD. Unpaired t-test. N=5 mice per group with 4 technical replicates, except 5xFAD for IL-6 N=4 mice.

| Cytokine | Cytokine concentration (pg/ml) | | Unpaired t-test | |
|---|---|---|---|---|
| | WT 11-12 mo (N=5) | 5xFAD 11-12 mo (N=5) | p-value | Significance |
| IL-6 | 6.3 ± 2.3 | 5.7 ± 0.3 | 0,6944 | ns |
| IL-10 | 20.1 ± 5.2 | 4.1 ± 1.1 | 0,0003 | *** |
| MCP-1 | 0.2 ± 0.03 | 0.2 ± 0.06 | 0,6526 | ns |
| IFN-γ | 2251.9 ± 653.4 | 923.5 ± 332.5 | 0,0067 | ** |
| TNF | 3.4 ± 0.3 | 3.6 ± 1.0 | 0,7268 | ns |
| IL-12p70 | 1192.5 ± 297.3 | 794.0 ± 425.1 | 0,163 | ns |
